# Striatal dopamine encodes the relationship between actions and reward

**DOI:** 10.1101/2022.01.31.478585

**Authors:** G. Hart, T.J. Burton, C.R. Nolan, B.W. Balleine

## Abstract

Although the role of striatal dopamine in Pavlovian conditioning and in habits has been reasonably well described, relatively little is known about its function in goal-directed action. In this study we trained hungry rats on two lever press actions for distinct food outcomes and recorded dopamine release in the dorsomedial striatum as these action-outcome associations were encoded and subsequently degraded. During initial training the lever press actions generated bilateral dopamine release that was found to reflect the predicted action value. This value was updated by the prediction error generated by the feedback produced by contact with the outcome, or its absence, after the press. Importantly, hemispheric dopamine release became increasingly lateralized across the course of training, with greater release in the hemisphere contralateral to the press. Using video analysis and multiple different measures, we could find no evidence that the degree of lateralized release was associated with movement; rather, we found that it tracked the strength of the action-outcome association, increasing and decreasing with increments and decrements in the contingency between specific actions and their consequences. Similar results emerged whether the rewards were delivered on ratio or interval schedules of reinforcement and whether we used unpaired outcome delivery or outcome-identity reversal to modify the specific contingencies. These findings suggest that, whereas moment-to-moment fluctuations in action value are reflected in bilateral dopamine release, a second signal broadcasts the overall strength of specific action-outcome relationships via the difference between contralateral and ipsilateral release during actions.

---

The capacity for goal-directed action is a core function that allows animals to encode the consequences or outcome of their actions to maintain adaptive behavior in a changing environment(1,2). Recent evidence suggests that action-outcome encoding depends on a prefronto-striatal circuit focused on the posterior dorsomedial striatum (pDMS)(3–6), with initial learning and the subsequent updating of these associations mediated by plasticity at two types of principal neuron(7): the striato-nigral direct spiny projection neurons (dSPNs), which express dopamine D1 receptors, and striato-pallidal indirect SPNs (iSPNs) expressing the D2 receptor(8) (9,10). Importantly, this plasticity appears to require the integration of glutamatergic inputs from cortical, thalamic and limbic regions with the input from midbrain dopaminergic neurons(11–18).

Although much has been learned about the role of striatal dopamine in the cellular and circuit level plasticity necessary for reflexive movements and skills (19–21) relatively little is known about its role in the acquisition and performance of goal-directed actions. Here we show, using a continuous, self-paced instrumental conditioning task, that, over the course of acquisition, goal-directed actions are associated with initially bilateral but increasingly hemispherically lateralized dopamine release that reflects the strength of specific action-outcome associations, tracking increments and decrements in the action-outcome contingency. In addition, trial-by-trial action values were reflected in a bilateral action value signal, reflecting action-related reward predictions that was itself adjusted by bilateral reward prediction errors occurring during exposure to the outcome (positively when presented, negatively when withheld). These findings suggest, therefore, that bilateral dopamine release during goal-directed action conveys both moment-to-moment adjustments in, and the long-term associative strength of, the action-outcome association.

## Dopamine release during the instrumental action is initially bilateral but becomes progressively more lateralized as training progresses

We utilized a genetically-encoded dopamine indicator, dLight1.1(22) (**Fig 1A, Fig S1A-B**), to measure rapid changes in dopamine release in the pDMS during the acquisition and performance of goal-directed actions. This indicator inserts circularly permutated GFP into dopamine D1 receptors such that the binding of dopamine causes an increase in fluorescence, which we measured using fiber photometry. We profiled dopamine release during goal-directed learning and subsequent performance, and then assessed effects specific to changes in the strength of the action-outcome association when this was modified using a contingency degradation treatment.

**Fig. 1.**
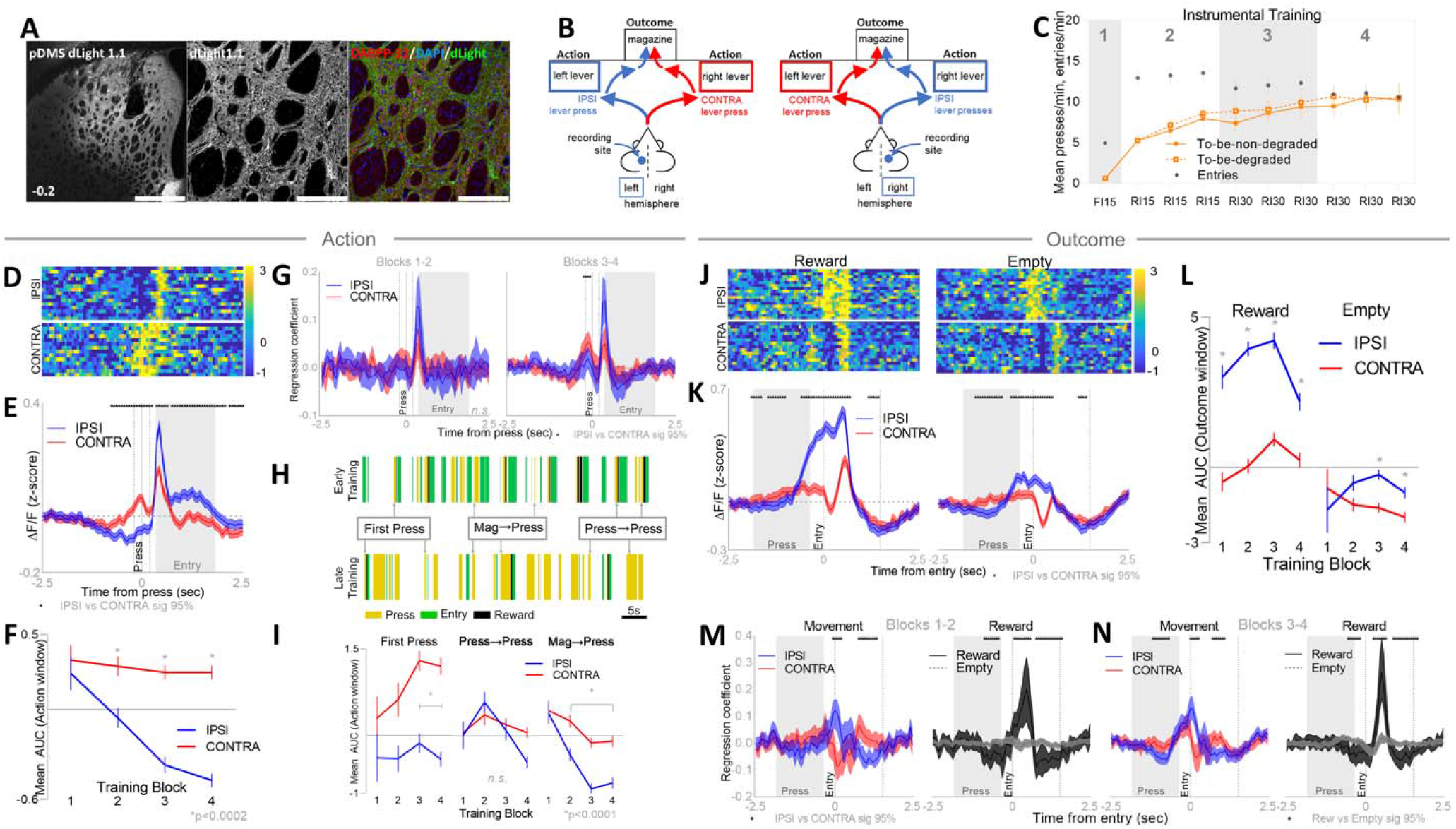
Dopamine release in the pDMS is lateralized during goal-directed actions and their outcomes. **(A)** Left: Location of fiber cannula and expression of dLight1.1 in the pDMS (0.2mm posterior to bregma), scale bar =1000μm. Center: from the same slice, dLight1.1 (grey). Right: same as center, dLight1.1 (green), DARPP-32 (red) and DAPI (blue), scale bar=200μm. **(B)** Rats were trained with alternatingly presented levers to the left and right of the reward magazine, designated as IPSI and CONTRA relative to the recording hemisphere. **(C)** Mean (±SEM) presses per minute on each lever and magazine entries per minute across each day of training. Numbers in grey specify training blocks. **(D)** Example trials showing z-scored ΔF/F dLight signals over time aligned to IPSI (top) and CONTRA (bottom) lever presses, x-axis indicated in panel E. **(E)** Mean (± 95% bCI) z-scored ΔF/F averaged across all IPSI (blue) and CONTRA (red) actions during instrumental training aligned to lever press (x=0), dotted lines either side indicate Action Window (±0.2s from press), grey shading indicates window within which the majority (>80%) of subsequent magazine entries occurred. Asterisks indicate a significant difference between IPSI and CONTRA signals at a given timepoint (95% bCI). **(F)** Mean (± SEM) area under the curve (AUC) of the z-scored ΔF/F signal during the Action Window for all IPSI and CONTRA lever presses, averaged within each training block. Asterisks indicate significant differences between IPSI and CONTRA. **(G)** Mean regression coefficients (± 95% bCI) for IPSI and CONTRA signals aligned to lever press for blocks 1-2 (left) and blocks 3-4 (right) of training, asterisks indicate significant IPSI vs CONTRA difference (95% bCI). **(H)** Representative sequence of events over 2 min for one rat during Block 1 of training (top) and the same rat in Block 4 of training (bottom), lever presses in yellow, entries in green and rewards in black, example of press types identified. **(I)** Mean (± SEM) AUC of the z-scored ΔF/F signal for IPSI and CONTRA lever presses, subdivided according to press type, averaged within each training block. Asterisks indicate significant IPSI vs CONTRA differences. **(J)** Example z-scored ΔF/F dLight signals over time aligned to rewarded (left) and non-rewarded (empty, right) magazine entries after IPSI (top) and CONTRA (bottom) presses, aligned to x-axes in panel K. **(K)** Mean (±95% bCI) z-scored ΔF/F averaged across all rewarded (left) and empty or non-rewarded (right) magazine entries after lever presses during instrumental training. Outcome Window (+1.5s after entry) indicated by dotted lines. Shading indicates window in which the majority (>80%) of preceding lever presses occurred. **(L)** Mean (± SEM) AUC of the z-scored ΔF/F signal during the Outcome Window for all rewarded (left) and empty (right) magazine entries after IPSI and CONTRA lever presses, averaged within each training block. Asterisks indicate significant IPSI vs CONTRA differences. **(M)** Mean regression coefficients (± 95% bCI) for dLight signals during blocks 1-2 of training, aligned to magazine entries after IPSI (blue) and CONTRA (red) lever presses (left), and aligned to rewarded (dark grey) and non-rewarded (empty, light grey) entries. Asterisks indicate significant differences between waveforms (95% bCI). **(N)** Same as **(M)** for blocks 3-4 of training.

Rats were trained on a goal-directed instrumental conditioning protocol in which two levers, located to the left and right of a central food magazine, each delivered distinct rewards (grain pellets and sucrose solution) into the magazine on increasing random interval (RI) schedules of reinforcement across sessions (**Fig 1B**). Response rates on each lever and mean magazine entry rates across days of training are presented in **Fig 1C**. Levers were grouped according to subsequent treatment as to-be-degraded or to-be-non-degraded, with side of lever (left and right) counterbalanced within each condition. Rats increased responding on both levers across days (significant linear trend F(1,20)=62.5, p=1.40e-7), and there were no differences in the response rates of each action (F’s<1.0).

Due to the lateralized nature of the task (actions performed to the left or right of the reward magazine), dopamine signals during lever presses (Actions) and subsequent magazine entries (Outcomes) were categorized hemispherically according to the side of the action relative to the dLight recording site: i.e., for animals with left hemisphere fiber photometry cannulae, left lever presses (and subsequent entries) were categorized as ipsilateral (IPSI) and right lever presses and their subsequent entries were categorized as contralateral (CONTRA; **Fig 1B**). To characterize the dynamics of dopamine release during all actions over task acquisition, we subdivided the training into 4 blocks (indicated in **Fig 1C**) and examined peri-event z-scored ΔF/F signals aligned to IPSI and CONTRA lever presses in a number of ways. IPSI and CONTRA presses were clearly distinct when examining waveforms at either the single event (**Fig 1D,** individual examples) or aggregate (**Fig 1E**, all presses) level. To analyze these waveform differences (**Fig 1E**), we applied a bootstrapped confidence intervals analysis procedure(23) (bCI; **Methods**) specifically designed for interrogating event-related transients in fiber photometry data. This analysis confirmed that pDMS dopamine release during an action was significantly greater in the hemisphere contralateral to the action compared to the ipsilateral hemisphere (**Fig 1E,** CONTRA>IPSI, 95% bCI). This difference was observed prior to, during and shortly after lever press initiation, with the relationship inverting shortly after the press (IPSI>CONTRA, 95% bCI), when animals turned in the opposite direction to enter the magazine.

To assess how this striking pattern of lateralized dopamine release behaved over the course of training, we calculated the area under the curve (AUC; trapezoidal method) during a window of time surrounding the lever press action (0.2s before to 0.2s after the lever press). This window was determined by examining the temporal dynamics of all lever press-magazine entry sequences (**Fig S1C**), and constrained by the reward delivery, which commenced 0.2s after a rewarded press. Dopamine release during the Action window was stable for CONTRA presses over the course of training but was progressively suppressed (relative to baseline) during IPSI presses (**Fig 1F**). There was no detectable difference in release during IPSI and CONTRA presses in Block 1 (p=0.6), but significant differences emerged in Blocks 2-4 (k=6, p-critical=0.008, p’s < 0.0002), indicating that the hemispherically lateralized dopamine signal during Actions developed across the course of training.

To parse the signals specifically associated with presses from those related to subsequent magazine entries, we fitted a linear (multiple) regression model(24) (see **Methods**), which isolated the variance in the dLight signal associated with IPSI and CONTRA lever presses, IPSI and CONTRA magazine entries (i.e. entries that followed IPSI and CONTRA presses), and rewarded and empty magazine entries. We modeled these signals across the first and second halves of training (Blocks 1-2 and Blocks 3-4), to compare the isolated IPSI and CONTRA effects during actions as training progressed. These modeled dLight signals during IPSI and CONTRA presses are presented as regression coefficients in **Fig 1G**. We applied the bCI analysis to statistically compare IPSI and CONTRA coefficients and confirmed that dopamine release during actions was comparable in both hemispheres in the first half of training, but that release was significantly greater in the hemisphere contralateral to the action during the later stages. This emergent effect was restricted to the period immediately preceding the Action within the Action window, **Fig 1G,** right, CONTRA>IPSI, 95% bCI).

It is possible that the progressively lateralized dopamine response reflects the initiation of action sequences rather than their ongoing performance(25) and so we examined whether organizing lever presses by their relationship to other events (i.e., other lever presses and magazine entries) yielded distinct profiles of dopamine release. **Fig 1H** illustrates a subset of behavioral sequences for one rat both early and late in training. As training progressed, presses and entries became loosely clustered into press-press and press-entry sequences interspersed with isolated lever presses. We therefore divided lever presses into three categories according to the immediately preceded event: (1) First Presses, for which neither a lever press nor an entry had occurred in the preceding window; (2) Press→Presses which were lever presses that followed another press in an action sequence; and (3) Mag→Presses, which were lever presses that followed a magazine entry. The total number of each press type in each training block is presented in **Fig S1D**. The mean AUCs during the Action window for each lever press category over the course of training are presented in **Fig 1I.** When the data were collapsed across training, dopamine release was significantly lateralized during First presses (k=15, p-critical=0.003, IPSI vs CONTRA p=1.61E-25) and Mag→Presses (IPSI vs CONTRA p=1.75E-17), but not Press→Presses (IPSI vs CONTRA p=0.02). This difference developed as training progressed; there were no differences between IPSI and CONTRA presses of any type in Block 1 (p’s >0.2), but a significant difference across Blocks 3-4 for First Presses (p’s<2.80E-09, Block 2 p=0.006) and across Blocks 2-4 for Mag→Presses (p’s<0.0003). Therefore, the emergent lateralization of dopamine release during actions over the course of training was associated specifically with the initiation of action sequences (First Presses) and those actions that directly followed a magazine entry in sequence (Mag→Presses). To broadly assess the relationship between dopamine release lateralization and response vigor, we calculated the correlation between the session-averaged dopamine release (AUC during the Action window) and the session-averaged inter-press interval **(Fig S1E)** as well as the correlation between individual lever press AUCs and the inter-event interval (the interval between each press and the immediately preceding press or entry, **S1F**). We found no significant evidence in these analyses to explain the emergent lateralization of dopamine release over the course of training (see the caption of **Fig S1** for further details).

These results show, therefore, that: (i) pDMS dopamine release became hemispherically lateralized during the acquisition of goal-directed actions; (ii) was specific to the direction of the action, with relatively higher release in the hemisphere contralateral vs. ipsilateral to the response direction; and (iii) was not obviously related to action speed or vigor.

## Dopamine release during retrieval of the instrumental outcome reflects both movement and reward value

Having characterized the pDMS dopamine release profile during the instrumental action component of goal-directed learning and performance, we next sought to profile dLight signals during outcome retrieval. Outcomes were defined as magazine entries that followed Actions (lever presses) and were categorized as being either Rewarded (rewarding outcome present) or Empty (no rewarding outcome present). Example trials showing the z-scored ΔF/F dopamine signals aligned to rewarded and empty magazine entries that followed IPSI and CONTRA lever presses are presented in **Fig 1J**. The peri-event waveforms for each Outcome category averaged across all animals during instrumental training are displayed in **Fig 1K**. Distinct dopamine release profiles were observed for magazine entries that followed IPSI vs. CONTRA presses, and also for entries that were rewarded or not rewarded. bCI analyses on the z-scored ΔF/F dopamine signals (**Fig 1K**) confirmed that dopamine release prior to the magazine entry reflected the lateralization of the preceding Action (CONTRA>IPSI), which inverted (IPSI>CONTRA) as animals approached and entered the magazine. The lateralized signal immediately preceding magazine entry appeared to reflect the direction of the animal’s movement: following an IPSI press, animals made a contralateral turn to enter the magazine (and vice versa for CONTRA presses). Additionally, the magnitude of the dopamine signal appeared diminished for empty relative to rewarded entries (**Fig 1K** left vs. right).

To assess Outcome-related dopamine release as training progressed, we calculated AUCs in the Outcome window, defined as 0-1.5s after magazine entry onset. This window was chosen to capture the entry period for the majority of animals (**Fig S1C**). The mean AUCs (**Fig 1L**) confirmed that rewarded magazine entries were associated with higher dopamine release than non-rewarded empty entries across training (k=32 p-critical =0.0015, IPSI→Rew vs. IPSI→Empty Blocks 1-4, p’s<5.8E-05; CONTRA-Rew vs CONTRA-Empty Blocks 2-4 p’s<0.0002). Further, greater dopamine release was observed during magazine entries that followed IPSI lever presses compared to those after CONTRA presses (IPSI→Rew vs. CONTRA→Rew Blocks 1-4 p’s <8.6E-07, IPSI→Empty vs. CONTRA→Empty Blocks 3-4 p’s<0.0012, note there are very few empty magazine entries in Block 1). Taken together, these results indicate an effect of both reward and movement on Outcome-related dopamine signals. However, unlike release during lever press actions, the Outcome-related movement signals were clearly and significantly lateralized (IPSI>CONTRA) from the beginning of training (**Fig 1L**).

To disentangle the relative influence of movement and reward on the dLight signal, we entered each of these factors as separate coefficients into the multiple regression. The isolated signals associated with movement (IPSI vs. CONTRA) and reward (Reward vs. Empty), across the first and second half of training are presented in **Fig 1M-N**. Across both the first (**Fig 1M**) and second (**Fig 1N**) halves of training, movement signals (magazine entries following IPSI and CONTRA presses, left panels) were similar. There was significantly greater dopamine release associated with IPSI entries (**Fig 1M-N,** IPSI>CONTRA at the time of entry, 95% bCI), which subsequently inverted (CONTRA>IPSI, 95% bCI), relating to the direction of movement as the animals returned to the lever after exiting the magazine. Similarly, in the period prior to the magazine entry, when the lever press was performed, a lateralized signal was observed consistent with that observed in the Action window and, importantly, was only statistically significant in the second half of training (**Fig 1N,** left, CONTRA>IPSI, 95% bCI). There was also a pronounced reward component to the magazine entry aligned signal across all of training (**Fig1 M-N,** right panels), wherein rewards were accompanied by a significant spike in dopamine release after entering the magazine (i.e., during consumption; Reward>Empty, 95% bCI).

## Bilateral dopamine release during the action reports the updated action value

Our results reveal that during instrumental training, dopamine release in the pDMS is consistently greater in the hemisphere contralateral to the action. However, the evolution of this signal across training differs in each component of the action→outcome sequence: dopamine release progressively diverges between the ipsilateral and contralateral hemispheres during the lever press action, whereas during outcome retrieval it is lateralized from the outset and remains largely unchanged as training progresses. As such, these results suggest that changes in dopamine release across training provide a signal to update the value of the action for goal-directed learning rather than the value of the outcome. To assess this hypothesis more directly, we next sought to examine whether dopamine release during goal-directed actions is modulated by a reward prediction error generated within the Action and Outcome windows described above.

We calculated reward prediction-errors for each action-outcome association (IPSI and CONTRA) across training for each animal, and used these to update action values (V) using a simplified reinforcement learning rule(26). According to this model, at the time of magazine entry (at time t) a prediction error is calculated, which is used to update the subsequent action value (at time t+1) according to: V(t+1)=V(t)+ δ(t) (**Fig 2A**), where δ is the prediction error modulated by a learning rate parameter (α) according to: δ(t)=α[R(t)– V(t)], where R is the reward value, set to 0 for no reward or 1 for reward, to generate a negative δ on non-rewarded entries and a positive δ on rewarded entries. Importantly, this simplified model assumes that reward predictions are only updated by experience with the consequences of the action (i.e., a rewarded or empty magazine entry). Therefore, during lever press sequences in which the rat doesn’t enter the magazine V will remain unchanged.

**Fig. 2.**
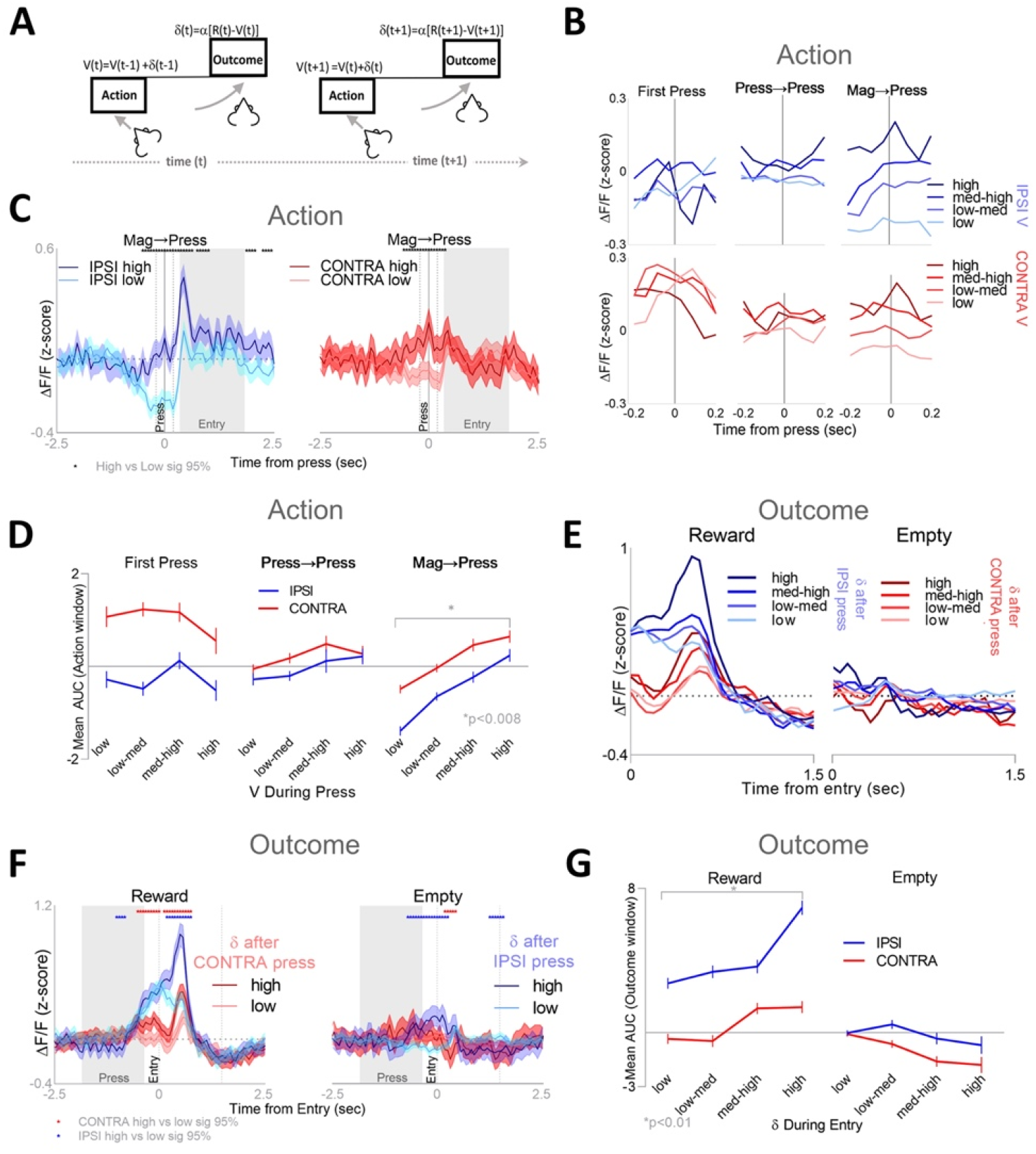
Dopamine reflects action values and outcome reward prediction errors. **(A)** Action values are updated by summing the previous action value with the reward prediction error calculated during the preceding magazine entry (Outcome). **(B)** Mean z-scored ΔF/F signals during the Action window (−0.2s before press to 0.2s after press) for IPSI (blues) and CONTRA (reds) lever press subtypes, subdivided according to action value (V). **(C)** Mean (±95% bCI) z-scored ΔF/F signals aligned to IPSI (left) and CONTRA (right) Mag→Presses with high (dark colors) or low (light colors) V-values. Asterisks indicate significant high vs low difference (95% bCI). **(D)** Mean AUC (±SEM) for signals shown in (B). Asterisk indicates significant high vs low difference for both IPSI and CONTRA. **(E)** Mean z-scored ΔF/F signals aligned to the Outcome window (0s to 1.5s after entry) for rewarded (left) and empty (right) magazine entries after IPSI (blues) and CONTRA (reds) lever press subtypes, subdivided according to reward prediction error (δ) calculated during that same entry. **(F)** Mean (±95% bCI) z-scored ΔF/F signals aligned to rewarded (left) and non-rewarded (empty, right) magazine entries with high and low prediction error. Red asterisks indicate significant (95% bCI) high vs low difference for CONTRA entries (i.e., entries after a CONTRA press) and blue for IPSI entries **(G)** Mean AUC (±SEM) for signals shown in (E). Asterisk indicates significant high vs low difference for IPSI and CONTRA.

**Fig 2B** shows the mean z-scored ΔF/F signals during the Action window across a range of V values (low-high: 0-1) corresponding to the calculated action value (V) at the time of the action. Whereas there was limited evidence for dopamine modulation during First Presses or Press→Presses, dopamine release during presses that followed magazine entries (Mag→Presses) appeared to be modulated by V. We directly compared the waveforms for Mag→Press actions with high vs. low V values (**Fig 2C**) and confirmed that there was significantly greater dopamine release for both IPSI and CONTRA lever presses (high vs low bCI sig 95%). This relationship between dLight activity and action value was not observed for First Presses or Press→Presses (**S2A**). These findings were bolstered by examining the mean AUCs during the Action window across a range of V values for each press type (**Fig 2D**). There was a clear relationship between dopamine release and V for IPSI and CONTRA Mag→Presses (k=6, p­critical=0.008; V(low) vs. V(high): Mag→IPSI, p=2.21E-22, Mag→CONTRA p=2.79E-12), but not for First Presses or Press→Presses (V(low) vs. V(high): First Presses p’s>0.1; Press-Presses p’s > 0.02). Note that, as our model assumes that prediction error is only generated during magazine entries, this finding suggests that these changes in dopamine release during the Action-window reflect the updating of action values. However, to ensure that this assumption did not drive the observed pattern of results, we modified the model to allow V to be updated on every press. In this case, all non-rewarded presses resulted in negative δ (i.e., presses followed by empty magazine checks and presses followed by other presses), and only presses followed by rewarded entries accrued a positive δ. The mean AUCs during presses across binned values of V using this model are presented in **Fig S2B**. Although there was generally more variability, the findings remained the same: dopamine release during presses that had followed magazine entries was positively related to V (k=6, p-critical=0.008; V(low) vs. V(high): Mag→IPSI p=3.36E-08, Mag→CONTRA p=0.0008), and there was no significant relationship between dopamine release and V during other press subtypes (V(low) vs. V(high): First Presses and Press→Presses, p’s > 0.05).

## Dopamine release elicited by the instrumental outcome reports a reward prediction error

Our results confirm previous reports(27) that bilateral dopamine release in the pDMS during the lever press reflects action values, and we have extended this finding to the free-operant condition and show that action value representation is only discernable for updated action values following a magazine entry. This suggests that it is only after contact with reward (or non-reward) – and the consequent exposure to the prediction error – that action values are updated.

To further investigate this finding, we analyzed the magnitude of both reward prediction error (δ) and dopamine release during Outcome retrieval. Magazine entries were organized according to the magnitude of the error (δ) generated by the model (from high to low), assuming δ is only calculated during entries that follow lever presses. The mean z-scored ΔF/F signals during the Outcome window across differing δ values, which are positive for rewarded entries and negative for unrewarded entries, are presented in **Fig 2E.** During rewarded magazine entries (**Fig 2E, left**), there appeared to be a graded increase in dopamine release according to increasing δ, consistent with a reward prediction error signal. We assessed the relationship between δ and dopamine release statistically by comparing the waveforms of high vs. low reward prediction errors using the bCI procedure, shown in **Fig 2F.** We found evidence for reward prediction error signals in pDMS dopamine release after both Rewarded and Empty magazine entries; specifically, dopamine release was significantly greater during rewarded entries (after both IPSI and CONTRA lever presses) for the higher δ values (**Fig 2F** left, high>low, 95% bCI). There was also evidence that dopamine release during Empty magazine was significantly lower for the higher δ values (**Fig 2F** right, high<low, 95% bCI, note high error during empty magazine entries is negative). There was also some evidence during magazine approach for release associated with reward anticipation (**Fig 2F** (left-CONTRA, right-IPSI), high > low, 95% bCI).

Finally, we assessed the mean AUCs during the Outcome window across all binned δ values (**Fig 2G).** This analysis confirmed a positive relationship between dopamine release and positive reward prediction errors (δ) during the Outcome window when rewards were delivered (k=4 p-critical=0.01, δ(low) vs. δ(high): IPSI→Rew p=2.35E-17; CONTRA→Rew p=5.22E-05), and between negative reward prediction errors and unrewarded (Empty) magazine entries (δ(low) vs. δ(high): CONTRA→Empty p=0.0005). However, this latter relationship wasn’t significant for IPSI entries overall (IPSI→Empty, p=0.16), due to the inversion of the signal during the Outcome window (**Fig 2F,** right) in which the reward anticipation (high >low) persisted into the start of the outcome window, and then inverted towards the end (low >high). Generally, therefore, these results confirm that pDMS dopamine release during reward reports a positive prediction error and during non-reward a negative prediction error.

## Contingency degradation reduces response rate on the degraded action without causing systematic changes in motor movements

We have presented two clear effects of pDMS dopamine release at the time of the lever press action: bilateral moment-by-moment adjustments in action value signals and an emergent hemispheric lateralization that does not discernably reflect response vigor. While the former is consistent with previous findings, the progressive nature of the latter is novel and suggests, since recordings were taken from the striatal region at the heart of goal-directed control, that it may track the strength of the goal-directed action-outcome association. If true, then hemispheric lateralization of dopamine release should increase during acquisition and decline when the action-outcome contingency is degraded.

To investigate this possibility, we measured dopamine release during the degradation of one of the two action-outcome contingencies in each subject over the course of 10 degradation training sessions. Both levers continued to be presented in the same fashion and earned the same outcomes on the same interval schedule as in training, except that throughout each session rats received additional non-contingent (i.e., response independent) deliveries of one of the earned outcomes delivered on a random time (RT) schedule (**Fig 3A**). As a consequence, the non-contingent outcome was the same as the outcome earned on one lever - the ‘degraded’ lever - but was ‘different’ to that earned on the other ‘non-degraded’ lever. It has previously been demonstrated that, for goal-directed actions, response independent outcomes ‘degrade’ previously established associations between an action and its specific outcome, resulting in the selective reduction in performance of that action relative to actions which have not had this degradation manipulation(28–31).

**Figure 3.**
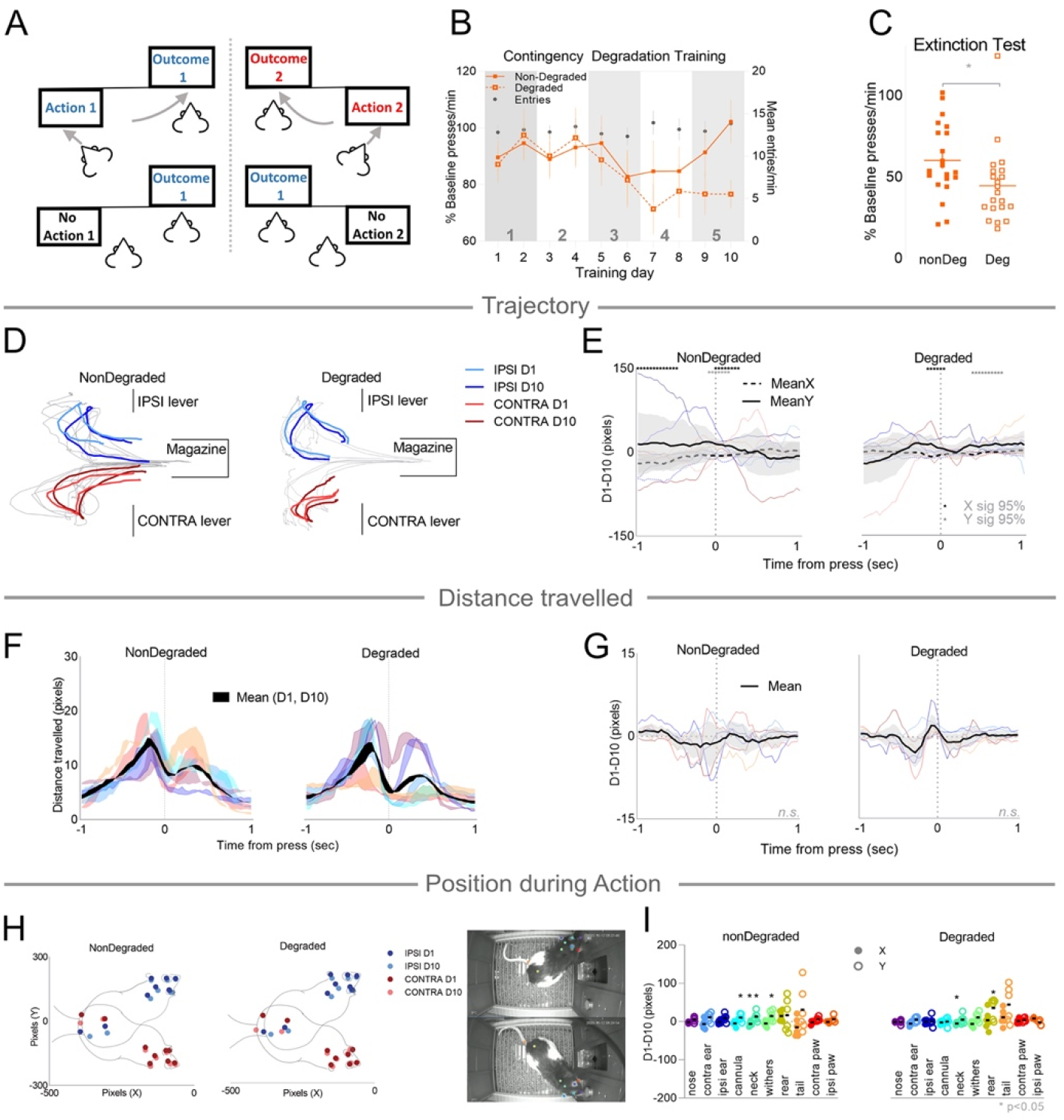
Tracking behavior and movement during contingency degradation. **(A)** In addition to each action-outcome pairing, rats received noncontingent (i.e., response-independent) deliveries of one of the earned outcomes. **(B)** Mean (±SEM) percent of baseline presses per minute on the non-Degraded (nonDeg) and Degraded (Deg) levers and mean (±SEM) magazine entries per minute (Entries) across 10 days of contingency degradation training. Numbers in grey indicate training blocks. **(C)** Percent of baseline presses per minute on the Degraded and non-Degraded levers for each animal during the extinction test. Lines indicate mean ±SEM. **(D)** The mean trajectory of the neck over a 2 second per-press period of all animals on each lever (colored lines), when it was either nonDegraded (left) or Degraded (right), for Day 1 (D1, light colors) and Day 10 (D10, dark colors) of contingency degradation training. Median trajectory for each individual animal shown in grey. **(E)** Difference scores (Day 1-Day 10) of peri-press co-ordinates in (D). X co-ordinates in dotted lines, Y co-ordinates in solid lines, the mean (±95% bCI) difference score of all animals is shown in black and the difference scores of each individual animal are in colored lines (blues for IPSI and reds for CONTRA). Lever press indicated by vertical dotted line. Asterisks indicate significant difference from zero for X co-ordinates (black) and Y co-ordinates (grey). **(F)** Mean distance travelled by the neck per frame across a 2 second peri-press period for each session (Day 1 and Day 10) on the nonDegraded (left) and Degraded (right) levers, with shading between the two lines to indicate the difference. The mean distance travelled for all animals is represented in black, with the median for each rat on each day plotted in the colored lines, (blues for IPSI and reds for CONTRA). Lever press indicated by vertical dotted line. **(G)** Difference scores (D1-D10) for distance travelled during the peri-press period represented in (F). Mean (±95% bCI) difference score of all animals is shown in black, and the difference scores for each individual animal are in colored lines. Lever press indicated by vertical dotted line. **(H)** The mean X-Y position of 10 tracked body parts (right, photos) for all rats at the time of press on the nonDegraded lever (left) and Degraded lever (right) on Day 1 (dark dots) and Day 10 (light dots) of training. **(I)** Difference scores for X (filled circles) and Y (open circles) co-ordinates for each tracked body part for each rat, colors matched to photo in (H), mean indicated by black bar. Asterisks indicate a significant difference between D1 and D10.

Lever press responding relative to baseline (Block 4 of instrumental training, see **Methods**) on the degraded and non-degraded levers across the 10 days of contingency degradation is presented in **Fig 3B**. There was no significant difference between responding on the degraded and nondegraded levers, averaged across all training days (F(1,20)=1.4, p=0.3), nor was there a significant linear trend across days, averaged across both levers (F(1,20)=2.2, p=0.2) There was, however, a significant lever x linear trend interaction (F(1,20)=5.4, p=0.03), indicating that responding significantly decreased on the degraded but not the nondegraded lever across training. The long-term effect of contingency degradation on instrumental responding was assessed the following day in a choice test in which both levers were presented simultaneously under extinction conditions (**Fig 3C**). Despite no rewards being delivered during this test, the suppression of responding on the degraded lever was maintained, with rats pressing the degraded lever significantly less than the non-degraded lever (F(1,20)=9.1, p=0.007), confirming the long-term effect of contingency degradation on action-outcome learning.

We assessed whether the divergence in response rate between degraded and non-degraded actions had any discernible effect on movement kinematics at the time of the lever press. To this end, we used Deeplabcut(32) with video footage to track a number of body parts on a subset of rats (n=6) during the first and last session (Day 1 and Day 10) of contingency degradation training. This allowed us to assess changes in the animals’ 2-D spatial trajectory, distance travelled around the lever press and body position during lever presses across training. These tracking data are shown in **Figure 3D-I**. To quantify the spatial trajectory of the lever press, we tracked the frame-by-frame movement of the marker aligned to the neck (the most reliably tracked midline point) through X-Y space across a 2-s period centered on the lever press (i.e. 1 sec before press until 1 sec after press) for all lever presses. The mean trajectory of all animals on each lever on Day 1 and Day 10 is presented in **Fig 3D**. Animals developed their own trajectory on approach to the lever and subsequent retreat from the lever (**Fig 3D**; individual median trajectories in grey), which appeared to remain relatively consistent from Day 1 of training to Day 10 on both Degraded and nonDegraded levers. To quantify the difference between Day 1 and Day 10, we calculated a difference score for each individual animal, as the median X and Y co-ordinates for each video frame on Day 1 minus Day 10, giving a difference score along each dimension across the 2-sec period. Individual and mean difference scores are presented in **Fig 3E**. Presses on both levers showed moderate shifts in trajectory along both X and Y axes (significance from zero, 95% bCI). Importantly, however, there was no evidence of a greater shift in trajectory on the degraded lever relative to the nondegraded lever on either dimension (X-Deg vs X-nonDeg and Y-Deg vs Y-nonDeg, 95% bCI, n.s.).

Next, we assessed distance travelled in the time preceding and succeeding the lever press, determined by the distance (pixels) the neck marker moved per video frame, represented in **Fig 3F**. The median distance travelled per frame for each rat on each day is plotted in the uniquely coloured lines, with shading between the two lines indicating the magnitude of the difference between Day 1 and Day 10 (mean for all animals in black). For both degraded and non-degraded lever presses, there was a consistent pattern in which distance travelled per frame increased immediately prior to the press and then decreased at the time of the press itself. We quantified the change in distance travelled from Day 1 to Day 10 by calculating difference scores (median distance travelled per frame on Day 1 minus Day 10), presented in **Fig 3G**. Although there were moderate shifts in distance travelled, these did not fulfil criteria for significance (difference from zero, 95% bCI n.s.).

Finally, we assessed the positioning of each rat during each lever press. To do this, we used 10 markers distributed across the body (**Fig 3H, right**). We extracted the XY co-ordinates of each marker during every press for every rat and calculated a median position in 2-D space for each of these markers for each rat. The mean spatial position for all rats on each lever is presented in **Fig 3H**. There was strong consistency in the positioning of animals from Day 1 to Day 10. The difference scores of the median X and Y co-ordinates for each rat between Day 1 and Day 10 for each marker are shown in **Fig 3I**; there was a significant shift in position of the cannula (Y), neck (X, Y) and withers (Y) on the non-degraded lever, and on the neck (X) and rear (Y) on the degraded lever (two-tailed t-tests, p’s <0.05). All other body parts remained spatially stable across Day 1 and Day 10 during each lever press.

Therefore, although there were some moderate changes detectable in 2-D trajectory and position during lever presses across contingency degradation training, these changes were similar on both levers and there was no evidence of systematic changes in motor movement that were specific to the degraded lever. Furthermore, although there appeared to be minor changes in distance travelled around degraded lever presses (a proxy for velocity) over degradation training these did not reach statistical significance.

## Contingency degradation reverses Action-specific but not Outcome-specific hemispheric lateralization of dopamine release

### Action-specific dopamine release – contingency degradation

Having verified that the structure of the lever-press response was largely unperturbed by the contingency degradation training, we evaluated the effect of this manipulation of pDMS dopamine release. The mean AUCs over the Action window for IPSI and CONTRA presses that were degraded or non-degraded are presented across degradation training blocks in **Fig 4A.** An effect of contingency degradation on dopamine release was apparent during both IPSI and CONTRA lever presses: Degradation was accompanied by a relative increase in dopamine release during IPSI presses, and a relative reduction in dopamine release during CONTRA presses, and these effects emerged over the course of degradation training. Specifically, there was no difference in either CONTRA or IPSI dopamine release during degraded versus nondegraded lever presses during the first training block **(Fig 4A**; IPSI and CONTRA Deg vs. nonDeg, Block 1 p’s > 0.1), but a significant difference was observed across all subsequent blocks for IPSI presses (k=10, p-critical = 0.005, IPSI Deg vs. nonDeg, Blocks 2-5, p’s <0.0001). There was a significant effect of contingency degradation for CONTRA lever presses in Blocks 2 and 4 (CONTRA Deg vs. nonDeg, p’s <3.18E-05), but not for Blocks 3 and 5 (p’s >0,1). To assess the possible source of this variability, we quantified the proportion of each press type in each block (**Fig S3A**), which revealed that in Block 3 (CONTRA) the distribution of press types differed from other blocks: Block 3 (nondegraded) had the lowest percentage of First Presses, as well as the highest percentage of degraded Press→Presses. We analyzed the waveforms for each press type separately for the entirety of contingency degradation training **Fig 4B**. The effect of contingency degradation was to reverse the hemispheric lateralization observed during initial training for First Presses and Mag→Presses. Specifically, dopamine release was relatively greater during both non-degraded vs. degraded CONTRA lever presses and during degraded vs. non-degraded IPSI presses (95% bCI). Importantly, although there were other epochs within the waveforms in which the signals differed, it was only during the Action window that we saw a modulation in both IPSI and CONTRA signals.

**Fig. 4.**
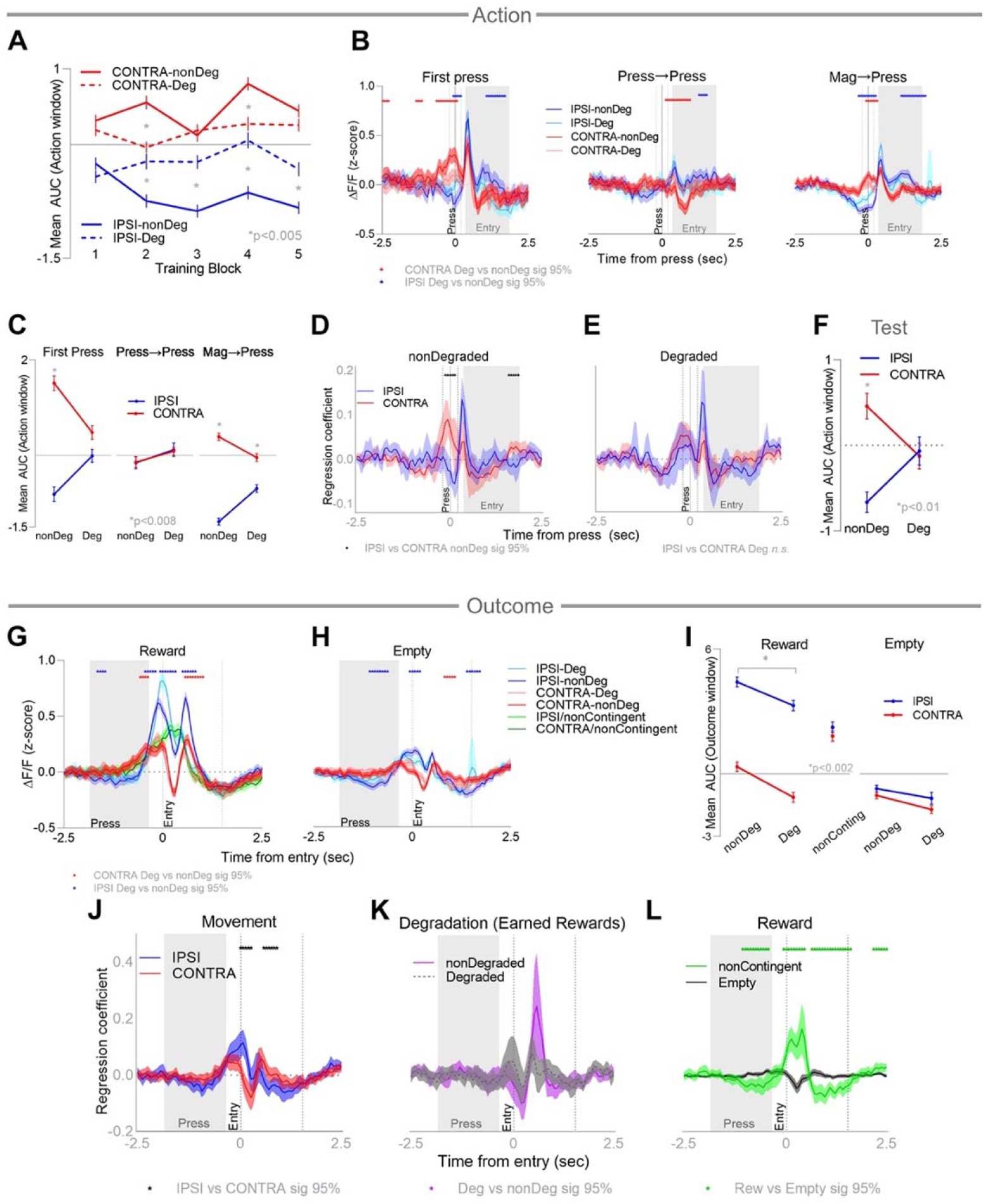
Dopamine lateralization during the Action, but not the Outcome, reflects instrumental contingency. **(A)** Mean (±SEM) AUC of the z-scored ΔF/F signal during the Action window for IPSI (blue) and CONTRA (red) lever presses that were Degraded or non-Degraded, averaged across training blocks of contingency degradation. Asterisks indicate significant Deg vs nonDeg differences. **(B)** Mean (±95% bCI) z-scored ΔF/F signals aligned to the lever press for IPSI (blues) and CONTRA (reds) press subtypes when they were Degraded (light colors) or non-Degraded (dark colors). Red asterisks indicate significant Deg vs nonDeg difference for CONTRA presses (95% bCI) and blue asterisks indicate the same for IPSI presses. **(C)** Mean (±SEM) AUC for z-scored ΔF/F signals in the Action window, averaged across training blocks 2-5. Asterisks indicate significant IPSI vs CONTRA differences. **(D-E)** Mean regression coefficients (±95% bCI) for dLight signals aligned to IPSI and CONTRA lever presses that were non-Degraded **(D)** and Degraded **(E)**, asterisks indicate significant IPSI vs CONTRA differences (95% bCI). **(F)** Mean (±SEM) AUC of the z-scored ΔF/F signal during the Action window for IPSI (blue) and CONTRA (red) lever presses that were Degraded or non-Degraded, averaged across the contingency degradation extinction test. Asterisks indicate significant IPSI vs CONTRA differences. **(G-H)** Mean (±95% bCI) z-scored ΔF/F signals aligned to the magazine entries that were rewarded **(G)** and empty **(H)** following IPSI (blues) and CONTRA (reds) lever presses, and during the retrieval of noncontingent rewards (greens), averaged across degradation training. Red asterisks indicate significant CONTRA Deg vs nonDeg differences, blue asterisks indicate the same for IPSI. **(I)** Mean (±SEM) AUC of the ΔF/F signal during the Outcome window, averaged across all of contingency degradation training. Left: Rewarded magazine entries that were contingent on IPSI (blue) or CONTRA (red) lever presses that were Degraded (Deg) or non-Degraded (nonDeg), or noncontingent rewards (nonConting). Right: Empty magazine entries following degraded or nondegraded IPSI or CONTRA lever presses. Asterisks indicate significant nonDegraded vs Degraded differences. **(J-L)** Mean regression coefficients (±95% bCI) for dLight signals aligned to magazine entries when they were IPSI vs CONTRA **(J)**, Degraded vs nonDegraded (earned rewards) **(K)** and Rewarded (noncontingent) vs nonrewarded (empty) **(L)**, asterisks indicate significant differences between waveforms (95% bCI).

These results were confirmed with the AUC analysis, which averaged the AUC during the Action window across the last 4 blocks of training (**Fig 4C**). Consistent with initial training, dopamine release was found to be greater during CONTRA relative to IPSI actions for nondegraded First Presses (k=6 p-critical=0.008; First Press IPSI vs CONTRA; nonDeg p=1.19E-25), importantly this effect was abolished for degraded First Presses (IPSI vs CONTRA Deg p>0.01). This same pattern of lateralization at the action was observed during nondegraded Mag→Presses (IPSI vs CONTRA nonDeg p=1.81E-64) and remained significant for degraded presses, albeit diminished (IPSI vs CONTRA; Deg p=1.54E-08). Consistent with the initial training data, there was no evidence of hemispheric lateralization in either degraded or non-degraded press sequences (Ipsi→IPSI vs. Contra→CONTRA, p’s>0.1).

We fitted a linear regression model to separate overlapping signals associated with lever presses (IPSI-Deg, IPSI-nonDeg, CONTRA-Deg and CONTRA-nonDeg), and magazine entries, compiled across all contingency degradation training. The isolated dLight signals during IPSI and CONTRA degraded and nondegraded lever presses are shown in **Fig 4D-E** (see also **S3B** for all presented on the same axis). Waveform analysis confirmed a significant CONTRA>IPSI difference during nondegraded lever presses (95% bCI) and this effect was indeed abolished when comparing IPSI and CONTRA degraded presses (IPSI vs CONTRA degraded, 95% bCI n.s.). Together, these results replicate our findings from training that dopamine signaling is lateralized during IPSI and CONTRA lever presses (specifically First Presses and Mag →Presses). However, and most importantly, this lateralization profile during the action was clearly dependent on the action-outcome contingency; degrading this contingency eliminated the lateralization. Therefore, hemispheric lateralization of pDMS dopamine release during goal-directed actions reflects more than just movement– it also scales with the strength of the action-outcome association.

### Action-specific dopamine release – degradation test

Finally, we measured dopamine release during the extinction test (**Fig 3C**). The total numbers of presses on the degraded and nondegraded levers across this test are shown in **Fig S3C**. The mean AUCs during the Action window are presented in **Fig 4F**. The pattern of results observed during the Action window across contingency degradation training was maintained on test: there was significantly greater dopamine release (AUC) during CONTRA lever presses relative to IPSI presses on the nondegraded lever (k=4, p-critical = 0.01, IPSI-nondeg vs CONTRA-nonDeg, p=3.95E-09). However, this hemispheric difference was abolished during degraded lever presses (IPSI-Deg vs. CONTRA-Deg p>0.7). As in training, the loss of lateralization during these actions was due to a bidirectional change in release: there was significantly less dopamine released during degraded than non-degraded CONTRA presses (CONTRA-nonDeg vs CONTRA-Deg p=0.007) and significantly more dopamine release during degraded than non-degraded IPSI presses (IPSI-nonDeg vs IPSI-Deg p=0.003). Therefore, the action-specific lateralization of dopamine release was significantly reversed when the instrumental action-outcome relationship was degraded, and this change was preserved under extinction conditions when no outcomes were delivered

### Outcome-specific dopamine release – contingency degradation

We also assessed dopamine release during the Outcome, presented in **Fig 4G-I**. The profiles of dopamine release during magazine entries that followed IPSI and CONTRA lever presses were similar under both nondegraded and degraded conditions (**Fig 4G-H**). However, their form differed for entries following noncontingent outcome deliveries (IPSI/noncontingent and CONTRA/noncontingent), which (by design) didn’t follow a lever press but were delivered at least one second after IPSI or CONTRA presses (**Fig 4G**). Waveform analysis indicated that there were several epochs in which dopamine release differed for degraded and non-degraded entries, however at no point did this shift indicate a loss of lateralization. The mean AUCs during the Outcome window for all contingency degradation training are presented in **Fig 4I**. There was no difference in dopamine release during magazine entries after the delivery of response-independent rewards during IPSI or CONTRA lever blocks (IPSI vs. CONTRA noncontingent, k=9, p-critical=0.006, p=0.13), although there appeared to be a general reduction in dopamine release during degraded compared to nondegraded magazine entries. For empty magazine entries this effect wasn’t significant (Deg vs. nonDeg: IPSI→Empty p=0.1; CONTRA→Empty, p=0.0064), however dopamine release was diminished during degraded compared to non-degraded reward retrieval (IPSI→Rew and CONTRA→Rew, Deg vs. nonDeg, p’s < 0.002), suggesting that reward-related dopamine release was depressed for degraded outcomes.

To further separate the effect of movement from reward and isolate the effect of contingency degradation on reward at the time of magazine entry, these factors were separately entered into the same linear regression model used to assess lever press signals. The regressed coefficients were, for both IPSI and CONTRA magazine entries: Rewarded-Degraded, Rewarded-nonDegraded, nonContingent and Empty, presented in **Fig 4J-L**. There was a significant effect of movement on dopamine signal during magazine entries (**Fig 4J,** IPSI>CONTRA, 95% bCI), which inverted towards the end of the magazine entry (CONTRA>IPSI 95% bCI) as rats left the magazine and turned back towards the lever. A separate regression analysis revealed there was no significant effect of degradation on these movement signals surrounding magazine entries (**Fig S3D,** IPSI and CONTRA Deg vs nonDeg 95% bCI n.s.), although there was a significant effect of movement (IPSI>CONTRA, 95% bCI) during both degraded and nondegraded magazine entries. Despite the indication in the AUC data for an effect of degradation on earned rewards (**Fig 4I**), this didn’t reach criteria for significance when examining the regressed waveforms (**Fig 4K**, 95% bCI n.s.). Finally, as expected, there was a significant effect of reward during the magazine entry, assessed by comparing the dLight signal during magazine entries after noncontingent reward delivery to that during empty entries (**Fig 4L**, nonContingent>Empty 95% bCI).

## Lateralization of pDMS dopamine release during the action is sensitive to the identity of the outcome associated with an action

Having established that the lateralization of pDMS dopamine release during goal-directed actions is sensitive to degradation of the contingency between the action and the outcome, we next sought to establish whether this phenomenon was likewise impacted by another change in contingency: that induced by a shift in outcome identity. Goal-directed action-outcome associations are underpinned by a relationship between the action and the specific sensory properties of the outcome. If dopamine lateralization during the lever press reflects the specific action-outcome association underlying goal-directed action, it should be apparent across different reinforcement schedules (i.e., across both ratio and interval schedules) and it should be sensitive to changes in outcome identity. To test this, we injected dLight1.1 into the pDMS of a naïve cohort of rats (**Fig S4A**) and trained them on the same goal-directed instrumental training protocol described, except the rewards were delivered on a random ratio (RR) schedule of reinforcement (**Fig 5A**). As should be expected, this training schedule resulted in the development of tighter chunks of lever press sequences, which generally terminated with reward delivery and entry to the magazine (**Fig S4B**). In contrast to interval schedule training, in which Mag→Presses and Press→Presses contributed similarly to the total presses, ratio training resulted in the dominance of Press→Press sequences (**Fig S4C-D**).

**Fig. 5.**
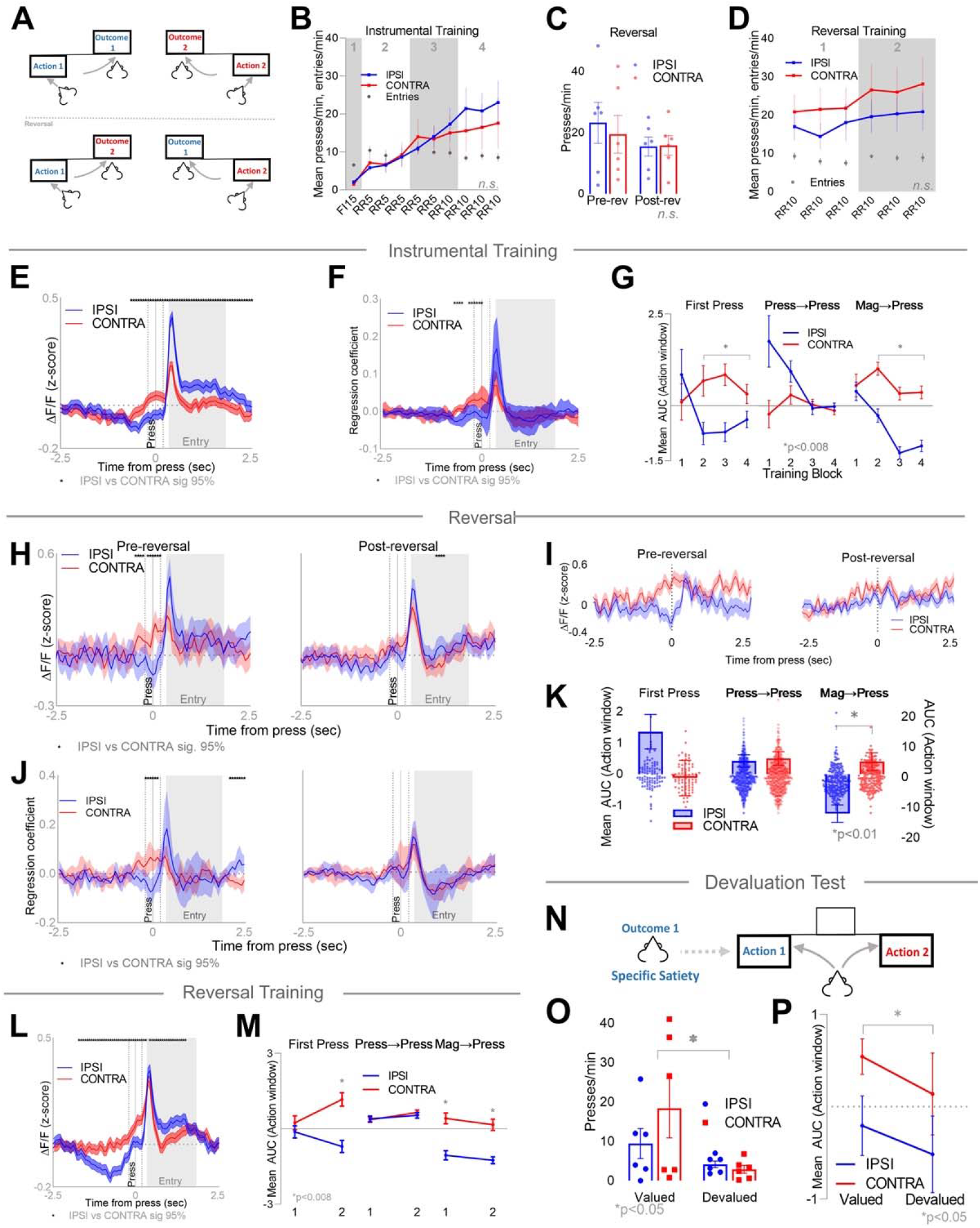
Dopamine lateralization is sensitive to outcome identity. **(A)** Rats were trained with two specific action-outcome pairings (instrumental training), the outcomes of which were then switched (reversal), followed by several days of training with the new action-outcome pairings (reversal training). **(B)** Mean (±SEM) presses per minute on the IPSI and CONTRA levers and mean (±SEM) magazine entries per minute (Entries) across 10 days of instrumental training. Numbers in grey indicate training blocks. **(C)** Presses per minute (open bars indicate mean ±SEM) for each rat on the IPSI (blue) and CONTRA (red) lever before (left bars) and after (right bars) the mid-session reversal. **(D)** Mean (±SEM) presses per minute on the IPSI and CONTRA levers and mean (±SEM) magazine entries per minute (Entries) across 6 days of reversal training. Numbers in grey indicate training blocks. **(E)** Mean (±95% bCI) z-scored ΔF/F signals aligned to IPSI and CONTRA lever presses, averaged for all instrumental training. Asterisks indicate significant IPSI vs CONTRA differences (95% bCI). **(F)** Mean regression coefficients (±95% bCI) for dLight signals aligned to IPSI and CONTRA lever presses, asterisks indicate significant IPSI vs CONTRA differences (95% bCI). **(G)** Mean (±SEM) AUC of the z-scored ΔF/F signals during the Action window for IPSI (blue) and CONTRA (red) lever press subtypes, averaged within each block of instrumental training. Asterisks indicate significant IPSI vs CONTRA differences **(H)** Mean (±95% bCI) z-scored ΔF/F signals aligned to IPSI and CONTRA lever presses pre-reversal (left) and post-reversal (right) for the reversal session. Asterisks indicate significant IPSI vs CONTRA differences (95% bCI). **(I)** Individual subject example from (H). **(J)** Mean regression coefficients (±95% bCI) for dLight signals aligned to IPSI and CONTRA lever presses pre-reversal (left) and post-reversal (right) for the reversal session. Asterisks indicate significant IPSI vs CONTRA difference (95% bCI). **(K)** Mean (±SEM) AUC of the ΔF/F signal (open bars, left y-axis) and AUC for each individual lever press (dots, right y-axis) during the Action window for press subtypes averaged across the reversal session after the reversal. Asterisks indicate significant IPSI vs CONTRA differences. **(L)** Mean (±95% bCI) z-scored ΔF/F signals aligned to IPSI and CONTRA lever presses averaged across all of reversal training. Asterisks indicate significant IPSI vs CONTRA differences (95% bCI). **(M)** Mean (±SEM) AUC of the ΔF/F signals during the Action window for IPSI (blue) and CONTRA (red) lever press subtypes, averaged within each block of reversal training. Asterisks indicate significant IPSI vs CONTRA differences. **(N)** One outcome was devalued by allowing rats to feed on it to satiety, immediately followed by a choice test with two levers available under extinction conditions. **(O)** Presses per minute on the IPSI (blue) and CONTRA (red) levers for each rat when they were valued (left) and devalued (right). Bars indicate mean ±SEM. Asterisk indicates significant valued vs devalued difference **(P)** Mean (±SEM) AUC of the z-scored ΔF/F signal during the Action window for IPSI and CONTRA lever presses when they were valued (left) and devalued (right). Asterisk indicates a significant valued vs. devalued difference.

Rats underwent initial instrumental training (**Fig 5B**) in which each lever earned distinct outcomes (grain pellets and sucrose solution) on increasing ratio schedules, trained in the same alternating fashion as the previous experiment. Response rates increased across days (**Fig 5B**, linear trend F(1,5)=62.05, p=0.001) and did not differ between IPSI and CONTRA levers (F < 1.0). In the second phase of the experiment, the outcomes associated with each response were reversed mid-session and delivered on the same reinforcement schedule (**Fig 5C**). There were no differences between IPSI and CONTRA press rates either pre-reversal or post-reversal (**Fig 5C**, F’s < 1.0). Training then continued for another 6 sessions on these reversed contingencies (**Fig 5D**) during which there was a significant increase in lever press rates across days (linear trend F(1,5)=8.4, p=0.03), but, again, no difference in IPSI and CONTRA press rates (F < 1.0).

The mean dLight signals surrounding IPSI and CONTRA lever presses, averaged across all instrumental training, are presented in **Fig 5E**. As in the prior experiment, we observed a significant CONTRA>IPSI difference leading up to, and during the lever press (CONTRA > IPSI, 95% bCI), which inverted after the press (IPSI>CONTRA 95% bCI). To isolate the signal associated with IPSI and CONTRA lever presses from that related to magazine entries, we used a linear regression model to separate IPSI and CONTRA presses and Rewarded and Empty entries, with these isolated press signals presented in **Fig 5F**, confirming a significant CONTRA>IPSI difference during the action window (CONTRA > IPSI, 95% bCI).

We then assessed the dLight signal during the Action window (AUC) for each lever press type separately across training (**Fig 5G**). As in the prior experiment, hemispheric lateralization of pDMS dopamine release was observed in First Presses and Mag→Presses; (k=6, p­critical=0.008, IPSI vs CONTRA averaged across Blocks 1-4; First Press p=2.55E-05, Mag→Press p=3.11E-20), but not Press→Presses (p=0.3). Importantly, and replicating the first experiment, this hemispheric lateralization of dopamine release developed across training; there were no statistically significant differences in dopamine release during the action between IPSI and CONTRA lever presses of any type in Block 1 (p’s>0.03). As before, we found no significant correlation between dopamine release during IPSI and CONTRA presses and either (i) the session-based lever press rate or (ii) the latency to press after a lever press or entry (**Fig S4E-F**), indicating that response vigor using these metrics cannot explain the hemispheric lateralization of pDMS dopamine release.

In the critical reversal phase, the outcomes associated with each action were reversed. The mean ΔF/F signals averaged across all IPSI and CONTRA lever presses performed pre- and post-reversal in this key session are presented in **Fig 5H** (with all presses averaged from one example rat shown in **Fig 5I**). Consistent with instrumental training, there was a significant CONTRA>IPSI difference in dopamine release during the lever press pre-reversal (**Fig 5H,** left, CONTRA>IPSI, bCI 95%). Importantly, however, this difference was abolished by outcome identity reversal (**Fig 5H,** right, IPSI vs CONTRA, bCI non sig.).

To confirm that this effect was specific to the lever press response, we isolated the signal associated with IPSI and CONTRA lever presses both pre- and post-reversal from that associated with subsequent magazine entries by applying the same linear regression model used in initial training. The regressed press-associated signals presented in **Fig 5J**. Confirming the prior finding, this analysis again found significant lateralization of dopamine release prior to reversal (**Fig 5J,** left, CONTRA>IPSI, bCI 95%), which was abolished post-reversal (**Fig 5J**, right). We calculated the mean AUCs for each press type post-reversal (**Fig 5K**) and found that pDMS dopamine lateralization was abolished during First Presses (IPSI vs CONTRA First Press, p=0.8), was preserved during Mag→Presses (IPSI vs CONTRA, k=3, p-critical=0.017, p=3.66E-05), and there remained no hemispheric difference during Press→Press sequences (p=0.07). Using Deeplabcut, we again conducted video analysis of the 2-D spatial trajectory, distance travelled and position during the peri-action period for each lever press before and after reversal, presented in **Fig S5**. We found no significant change in any of these movement metrics around the lever presses before or after reversal. Taken together, these results indicate that outcome identity reversal abolished hemispheric lateralization of pDMS dopamine release during lever pressing despite no obvious change in movement around the action.

In the third phase of the experiment, we sought to assess whether hemispheric lateralization of pDMS dopamine release would re-emerge with further training on the reversed contingencies as animals learned the new action-outcome associations. The mean ΔF/F signals averaged across reversal training for IPSI and CONTRA lever presses are presented in **Fig 5L**. The CONTRA>IPSI difference in dopamine release re-emerged during this training, however this hemispheric lateralisation preceded the press onset by longer than it did during initial training (CONTRA>IPSI, 95% bCI), and then inverted after the press (IPSI>CONTRA 95% bCI). The dopamine AUCs during each press type across blocks of reversal training are presented in **Fig 5M.** As in initial training, hemispheric lateralization during First Presses re-emerged over the course of reversal training. There was no hemispheric difference in dopamine release during training Block 1, but a significant difference was observed by Block 2 (k=6, p-critical = 0.008, First Press IPSI vs CONTRA; Block 1 p=0.3, Block 2 p=3.35E-07). As in initial training and reversal, there remained no difference in dopamine release during IPSI and CONTRA Press→Press sequences (p’s > 0.4), and the hemispheric lateralization during Mag→Presses persisted throughout reversal training (p’s < 2.68E-07). These data confirm, therefore, that continued training re-established the hemispheric lateralization observed during the initiation of action sequences.

Finally, we hypothesized that the re-emergence of the hemispheric lateralization of pDMS dopamine release during reversal training signaled that the new action-outcome contingencies had been acquired. In order to confirm this, we tested the rats for sensitivity to outcome devaluation using specific satiety(33). Rats were given free access to one of the earned outcomes for 30 min and were then given an extinction choice test with both levers available (**Fig 5N**). This was repeated on a subsequent day with the alternate outcome provided. The press rates on the valued and devalued IPSI and CONTRA levers during the devaluation tests are presented in **Fig 5O, Fig S4G**. Rats showed a clear devaluation effect in choice performance (i.e. valued > devalued lever presses) in accordance with the reversed contingencies (**Fig 5O;** main effect of value; Valued vs. Devalued F(1,5)=10.76, p=0.02), and there was no difference in overall response rates between IPSI and CONTRA lever presses (no main effect of side; IPSI vs. CONTRA, F < 1.0). Dopamine release during this test, presented as the mean AUC during the Action window (**Fig 5P**), reflected both the goal-directed action-outcome associations and the current outcome values: there was significant hemispheric lateralization (main effect of side; IPSI vs CONTRA, F(1,1040)=6.82, p=0.009), and a bilateral modulation according to the outcome value: dopamine release during the action was significantly lower when the outcome was devalued (main effect of value; Valued vs Devalued, F(,1040) IPSI=3.59, p=0.03, CONTRA=3.67, p=0.03). Together, these results confirm that dopamine lateralization during actions reflects the goal-directed action-outcome association and suggest that a shift in net (i.e., bilateral) dopamine reflects the devaluation-induced change in action value.

## Discussion

The decision to engage in an action to achieve a specific goal depends on integrating knowledge of the causal relationship between the action and its outcome with the current value of that outcome(3,6). Whereas the former involves encoding a relatively stable relationship across time, the latter is more dynamic and can fluctuate moment to moment with changes in motivational state and the immediate reward history of the action that alter action value. Here, we have shown that dopamine release in the pDMS during goal-directed action is sensitive to both sources of information, reflecting the overall strength of the specific action-outcome association and the current action value. We found that moment to moment fluctuations in bilateral dopamine release during the action were associated with adjustments to the action value signal in accordance with the reward prediction error during the immediately preceding outcome. However, we also found evidence of a generally more substantial and stable signal that emerged with experience and reflected the overall strength of the specific action-outcome relationships. This signal emerged in the difference between release in the hemisphere contralateral to the action and in the hemisphere ipsilateral to the action during action sequence initiation. Dopamine release reflects, therefore, both the strength of the action-outcome association and the current value of the action, the integration of which lies at the heart of the acquisition and performance of goal-directed actions.

Difficulty reconciling the function of striatal dopamine release in reward prediction, movement and motivation has provided a significant barrier to understanding its role in goal-directed action. A prevailing view has been that reward prediction errors are conveyed in the rapid ‘phasic’ firing of dopamine neurons, whereas slower variations in firing, as well as ‘tonic’ levels, support movement and motivation(34,35). There has, however, been little or no evidence for these slower or tonic signals using techniques that allow fast timescale measurements(36). Furthermore, while there is evidence for anatomically distinct dopaminergic axonal signaling related to locomotion and reward(25,37,38), it has remained a challenge to discern how they are dissociated at the level of dopamine release; i.e., parsing when increased dopamine release signals reward and when it signals action. Lateralized dopamine release during action performance has been interpreted as a movement signal(39–41), and indeed we saw some evidence of movement signals associated with reward retrieval responses. However, we were able to link the lateralized signal during lever press action sequence initiations directly to the strength of the action-outcome association and to show that this signal was distinct from the lateralized movement signals observed during subsequent reward approach and retrieval. Dopamine release was comparably elevated by ipsilateral and contralateral instrumental actions at the onset of training, consistent with bilateral encoding. However, this activity diverged over the course of training, with dopamine release maintained in the contralateral hemisphere while being reduced ipsilaterally. In contrast, release during reward retrieval was lateralized from the outset of training. Furthermore, whereas dopamine lateralization during reward retrieval persisted when the action-outcome relationship was degraded, lateralization was abolished during an action when the specific action-outcome association was degraded.

Importantly, this signal was not only sensitive to explicit degradation of the action-outcome contingency, it was also sensitive to changes in outcome identity. It is important to note that in the degradation and outcome switch manipulations the actions continued to be rewarded as in training and so the changes in release cannot reflect changes in reward value per se. Nor were these changes associated with any obvious or systematic changes in motor performance. Taken together with its replication across experiments and schedules of reinforcement, these findings provide, therefore, consistent evidence that this lateralized signal reflects a learning-induced change that tracks the strength of the specific action-outcome association necessary for the initiation of goal-directed action sequences. This finding is broadly consistent with the claim that SNc dopamine neuron activity is causally related to the initiation of lever press sequences(25). However, it is also clear that dopamine release can be rapidly shaped by local changes within the striatum proximal to the midbrain terminals, particularly those induced by other modulators, such as acetylcholine release from the cholinergic interneurons(7,42), it cannot be assumed, therefore, that the midbrain is the exclusive (or even primary) source of signal modulation.

Precisely how this hemispheric lateralization emerges to support the action-outcome association remains to be determined. One possibility is that these associations are first encoded bilaterally in pDMS(6,7) – perhaps reflecting the function of a dopamine-related, bilateral eligibility signal(43) – with release modulated hemispherically with experience and resulting in the inhibition of ipsilateral release by either local processes or via direct descending inhibition onto dopamine neurons(44). Either way, the opposing effects of high and low dopamine states on dSPNs and iSPNs(8) will ensure that contralateral release favors the performance of actions controlled by that hemisphere, with reduced ipsilateral release simultaneously inhibiting alternative actions(9,45). Although speculative, this model anticipates that hemispheric lateralization will be lost when bilateral plasticity is most likely required to adjust behavior: i.e., during initial learning, during degradation and after a switch in outcome identity.

In contrast to the large but relatively stable changes in lateralized dopamine release, rapid, moment-to-moment changes in bilateral release were associated with updating action values. This finding is supported by previous reports based on changes in GCaMP activity in DMS-projecting dopamine neurons(24,27), and resonates with similar signals in the ventral striatum encoding value and reward prediction-error for stimulus-outcome associations(36). Importantly, we found that the only the updated action value was reflected in the dopamine signal, on trials immediately after contact with reward or non-reward during outcome retrieval. There was substantial evidence to support this conclusion: We found no evidence for action value coding during press-press sequences but, more importantly, there was clear evidence of both positive and negative outcome-induced prediction-error coding by dopamine release during reward retrieval, confirming that a dopamine error signal was indeed generated at this time to allow the value of the action that immediately followed that retrieval to be updated. These results confirm the longstanding assumption that, as in Pavlovian conditioning, dopamine release during exposure to the instrumental outcome is bilaterally modulated by the reward prediction error and it is this error that updates the action values. In the final experiment we demonstrated that changes in outcome, and thus action, value with specific satiety-induced outcome devaluation also engendered a shift in bilateral dopamine release while maintaining the hemispheric lateralization signaling the action-outcome association. Following an outcome devaluation treatment, animals showed clear hemispheric lateralization of pDMS dopamine release in addition to a bilateral downward shift in this signal when animals were pressing the devalued lever, providing a clear example of how action-outcome and action value signals are integrated at the level of striatal dopamine release to support goal-directed action.

## Materials and Methods

### Animals

The experiments were conducted with healthy, experimentally naïve wild-type outbred Long-Evans rats aged between 8-16 weeks old prior to surgery (244-646g prior to surgery), obtained from the Randwick Rat Breeding Facility at the University of New South Wales. Rats were housed in transparent plastic boxes of 2-4 in a climate-controlled colony room and maintained on a 12 h light/dark cycle (lights on at 7:00 am). All experimental stages occurred during the light portion. Water and standard lab chow were continuously available prior to the start of the experiment. All experimental and surgical procedures were approved by the Animal Ethics Committee at the University of New South Wales and are in accordance with the guidelines set out by the American Psychological Association for the treatment of animals in research.

#### Exclusions and counterbalancing

##### Contingency Degradation

Subjects were 30 rats (13 female). All training and lever allocations were fully counterbalanced between males and females, left and right hemisphere recordings and within housing boxes with regards to order of lever and outcome presentation and identity of lever and outcome being degraded. Following histological assessment for placement of dLight1.1 and fiber optic in the pDMS, there were 9 rats excluded for misplaced cannulas or virus injections, leaving a total of 21 rats in the experiment (10 female), with 6 right pDMS recordings (3 ipsilateral lever degraded) and 15 left pDMS recordings (7 ipsilateral lever degraded).

##### Outcome Identity Reversal

Subjects were 8 rats (4 female). All training and lever allocations were fully counterbalanced as in the Contingency Degradation experiment except that all recordings were made in the left hemisphere. Two rats were excluded from the study due to misplaced fiber optic cannulae, leaving a total of 6 rats in the experiment (3 female).

### Surgery

Rats were anaesthetized with 3% inhalant isoflurane gas with oxygen, delivered at a rate of 0.5L/min throughout surgery. Anaesthetized rats were placed in a stereotaxic frame (Kopf), and an incision was made down the midline of the skull, and the scalp was retracted to expose the skull. A 1.0 uL glass Hamilton syringe was lowered into the brain for infusions of dLight1.1 (pAAV5-CAG-dLight1.1, AddGene #111067-AAV5), which was delivered at a total volume of 0.75uL (0.1uL/min), with the syringe left in place for an additional 3 min to allow for diffusion. A fiber optic cannula (400um, 6mm, Doric) was then implanted above the injection site, and secured in place with dental cement, attached to the skull with 3 jeweller’s screws. Following surgery, rats were injected with a prophylactic (0.4 mL) dose of 300 mg/kg procaine penicillin. Rats were given a minimum of 3 weeks of recovery time following surgery to allow sufficient viral expression.

Surgical co-ordinates were pre-determined from pilot studies and varied slightly between males and females, with dLight infused into males at the co-ordinates (mm from Bregma) A/P:-0.5, M/L:±2.6, D/V:-4.8 and for females A/P:-0.4, M/L:±2.55, D/V:-4.8, with fiber optic placement targeted 0.1mm dorsal to these injection sites.

### Apparatus

Training was conducted in two MED Associates operant chambers enclosed in sound- and light-attenuating cabinets. Each chamber was fitted with a pellet dispenser capable of delivering a 45 mg grain food pellet (Bioserve Biotechnologies), as well a pump fitted with a syringe outside the chamber, capable of delivering 0.2 mL of 20% sucrose solution (white sugar, Coles, Australia) diluted in water, each delivered to separate compartments of a recessed magazine inside the chamber.

The chambers also contained two retractable levers that could be inserted individually on the left and right sides of the magazine. Head entries into the magazine were detected via an infrared photobeam. Unless otherwise stated, the operant chambers were fully illuminated during all experimental stages by a 3W, 24V house light located on the upper edge of the wall opposite to the magazine. All training sessions were pre-programmed on a computer through the MED Associates software (Med-PC).

### Fiber Photometry

Fiber photometry recordings were conducted using a dedicated fiber photometry processor and software (RZ5P Processor, Synapse, Tucker-Davis Technologies, TDT), which were used to control LEDs (465nm excitation and 405nm isosbestic) via LED drivers with a Fluorescence Mini Cube, measured with Newport photoreceivers, all from Doric Lenses. Patch cords used for recordings (400um, Doric) were photobleached for a minimum of 45mins at the start of each recording day. Light was measured at the tip of the patch prior to recording sessions and maintained at 10-20uW (for 465) and 3-7uW (for 405). LEDs were modulated and demodulated via Synapse software at 331 Hz (465) and 211 Hz (405) and low-pass filtered (6Hz). MedPC signals were sent to RZ5P/Synapse, to indicate reward deliveries, magazine entries and lever presses, which were timestamped into the fiber photometry recordings.

### Behavioral Protocol and Food Restriction

#### Food restriction (all experiments)

Rats underwent 2-4 days of food restriction prior to the onset of magazine training and this continued throughout the duration of the experiment. During this time, they received 5 g each of chow daily in addition to 1 g each of grain pellets and 2hr/daily exposures to sucrose solution. Rats were then given 10-20 g of chow daily from the fourth day until the end of the experiment. Their weight was monitored to ensure it remained above 85% of their pre-surgery body weight at all times.

#### Magazine training (all experiments)

Rats were given two days of magazine training prior to the onset of instrumental conditioning, which consisted of 15 deliveries each of pellets and sucrose into the magazine, delivered intermingled (but non-overlapping) at random intervals of 60 s.

#### Contingency Degradation Experiment

##### Instrumental training

Rats were trained on an alternating lever paradigm, in which each lever came out for a total of 15 minutes or 10 outcomes, whichever came first, in alternating fashion, such that each lever was presented twice (total of 20 outcomes available per lever). On Day 1 of instrumental training (Block 1), rats were trained to press both levers on a fixed interval 15-sec schedule (FI15), where reinforcements were separated by a fixed interval of minimum 15 seconds, but every response spaced by more than 15 seconds was reinforced. This continued until rats reached criterion of earning 40 outcomes in a single session.

Over the following 3 days (Days 2-4 of instrumental training, Block 2), rats were trained on an RI15 sec schedule (presses were reinforced on average every 15 secs), on the same alternating levers paradigm, with outcome criterion increased to 30 of each outcome in each session (15 outcomes per lever presentation). Over the following 6 days (Days 5-10, Blocks 3-4), rats were trained on the same paradigm on an RI30 sec schedule (presses reinforced on average every 30 seconds).

##### Contingency degradation

Over the following 10 days (5 x 2-day blocks), rats were trained on a contingency degradation schedule, in which rewards were earned on the same RI30 schedule with alternating levers, with one outcome (pellets or sucrose) additionally delivered non-contingently on an RI45 sec schedule, explicitly unpaired with instrumental responses or magazine entries (no free outcomes could be delivered within 1 sec of a lever press or magazine entry), delivered during both lever presentations, but not during the 1min interval between levers. This schedule was chosen to approximately match the number of free and earned outcomes of the same type (rats received 31 free outcomes on average at the start of training and 36 on average by the end of training).

##### Test

On the final day after contingency degradation, rats were given a choice test under extinction (i.e., no outcomes were delivered) with both levers for 20 minutes. Rats were placed in the chambers, and after a 1 min interval, the house lights were illuminated and both levers came out for 20 mins, after which the session ended.

#### Outcome Identity Reversal Experiment

##### Instrumental Training

Lever press training was administered as in the Contingency Degradation experiment with the following differences: after achieving criterion on the FI15 schedule (Block 1; 15 minutes or 10 outcomes per lever presentation), rats were trained on a Random Ratio-5 (RR5) schedule for 5 days (Block 2) where outcomes were earned after an average of 5 responses (15 minutes or 15 outcomes per lever presentation for the remainder of the experiment), followed by 4 sessions of RR10 training (Block 3).

##### Mid-Session Reversal

Maintaining the RR10 schedule of reinforcement, animals underwent a single session where the outcomes paired with each action were reversed mid-session. This session had 6 alternating lever presentations (order counterbalanced) where the initial action-outcome pairings remained intact for the first 2 presentations (i.e. R1-O1, R2-O2) and were reversed for the remaining 4 (i.e. R1-O2, R2-O1).

##### Reversal Training

Rats were trained on the reversed contingencies (i.e. R1-O2, R2-O1) for a further 6 sessions (RR10) as in Instrumental Training.

##### Outcome Devaluation Testing

All rats received two 1-hour pre-exposure sessions to their individual devaluation chambers prior to the outcome devaluation tests. For outcome devaluation tests, rats were placed in their devaluation chamber for 30 minutes with *ad libitum* access to either grain pellets or sucrose solution (counterbalanced for pairing with lever relative to recording hemisphere). Rats were then immediately placed in the experimental chambers where they were given a 5-minute choice test in extinction with both levers available. This test was conducted twice for each rat (after devaluation of each outcome separately) with a session of re-training on the intervening day with the reversed contingencies maintained.

### Histology and Immunofluorescence Staining

Within a week after the final contingency degradation or devaluation tests, the rats were perfused with 4% paraformaldehyde in phosphate buffer (PB), and the brains sliced coronally on a vibratome at 40μm. A minimum of 6 sections each from the pDMS (from +1.2mm to −0.6mm A/P from Bregma) were stained for dLight1.1 and DARPP-32. Sections were rinsed 3 times for 10 min each in 0.1 M phosphate buffered saline (PBS), then submerged in a blocking solution of PBS with 0.5% Triton and 10% normal horse serum (NHS) for 2 hrs at room temperature. Sections were then submerged in 600 μL of rabbit anti-GFP (1:1000, Invitrogen) and mouse anti-DARPP-32 (1:1000, BD Biosciences) diluted in PBS with 0.2% Triton and 2% NHS for 48 hrs at 4 degrees C. Sections were then rinsed 3 times for 10 min in PBS, before being submerged in donkey-anti-rabbit Alexa 488 (1:1000, Invitrogen) and donkey-anti-mouse Alexa 546 (1:1000, Invitrogen), diluted in PBS with 0.2% Triton and 2% NHS for 2 hrs at room temperature. Sections were washed twice with PBS and twice with PB for 10 min each and mounted with Vectashield mounting medium with DAPI (Vector Laboratories).

Placement of dLight1.1 injections and fiber cannulas were imaged 24-48 hours later under a confocal microscope (BX16WI, Olympus) with a 10x objective using the boundaries defined by Paxinos & Watson (2016). Rats were excluded from the analysis if the placement of the fiberoptic cannula or virus extended anterior to +0.2mm from bregma in the DMS, or cannula placement was outside of the boundaries of the pDMS.

### Fiber Photometry Analysis

Fiber photometry data were analyzed in custom MATLAB scripts. To achieve a motion-artifact corrected dLight signal, the demodulated dLight signal (465nm) and isosbestic control signal (405nm) were extracted from the recording software (Synapse), and the isosbestic signal was fitted to the dLight signal using a least-squares linear fit model for each recording session. Change in fluorescence (ΔF/F) was calculated as (dLight signal – fitted isosbestic)/fitted isosbestic. Peri-event ΔF/F signals were extracted for left lever presses, right lever presses and magazine entries, across a 10s window (5s before and 5s after event onset) and downsampled to 15.8 samples per second (i.e., non-overlapping windows of 64 samples were averaged into a single value). The ΔF/F signal was normalized within these windows to the first 2.5s of each 10s window (baseline period), to give z-scores indicating the mean change from baseline (ΔF/F minus the mean baseline ΔF/F divided by the standard deviation of the baseline ΔF/F).

For waveform and regression analysis, we employed a bootstrapping confidence intervals procedure (bCI)(23) on ΔF/F dLight signals or regression coeficients, aligned to the lever press or magazine entry. This approach allows for the direct comparison of waveform profiles for two groups of signals (e.g., IPSI vs CONTRA). For a given event category, the population of Z-scored ΔF/F signals were randomly resampled with replacement for the same number of trials in that category and the mean waveform calculated. This was repeated 1000 times (i.e., 1000 bootstraps) to create a distribution of bootstrapped means. The differences between mean resampled waveforms (i.e. IPSI - CONTRA) were calculated at each timepoint, and 95% CIs were then calculated for each timepoint in the series (taking the 2.5 and 97.5 centiles from the ranked bootstrap distribution, expanded by √(n/(n – 1)) to correct for the narrowness bias). CIs that did not contain zero were identified, with at least four such consecutive timepoints constituting a significant difference (imposed to minimize the detection of spurious transients and corresponding to a consecutive threshold period of ∼0.25s). For analysis of trajectory and distance travelled, we employed the same approach and applying the same consecutive threshold period of ∼0.25s (six frames).

For event-related analysis, the area under the curve (AUC; calculated via the trapezoidal method for integral calculation; trapz function in MATLAB) was calculated for each event across event windows: Action: 0.2s before to 0.2s after press; and Outcome: 0 to 1.5s after magazine entry. These AUCs were averaged to give mean AUCs for each event type.

Multiple regression analysis followed Parker et al., 2016. Briefly, raw dLight signal (465 nm) was filtered with a high-pass (0.05 / 0.1 Hz stopband / passband) for baseline and drift correction, then divided by the pre-filtered mean to generate a ΔF/F. This signal was then z-scored by dividing by its standard deviation. The total signal was then modelled as the sum of event kernels (−2.5-2.5 s) convolved with respective event streams, where an event stream contained a 1 at the signal sample closest to the event, and a 0 at all other times. Across the relevant sessions and within each recording site / animal, the coefficients of the kernels were fit using the linear regression class from the scikit-learn Python package (sklearn.linear_model.LinearRegression). We ran three separate regressions, in each case fitting the coefficients for all the events in the respective analyses shown in figures 1, 3 and 5.

### Changes in action values

Action values were modelled using a simplified temporal difference learning rule, in which value, V, was updated according to an error term (δ) generated during magazine entry according to δ(t)=α[R(t)-V(t)], whereby α was a learning rate parameter and R was set at 1 for rewarded entries and 0 for nonrewarded entries. The action value of the next action, V(t+1) was updated according to the sum of the preceding action value and its error V(t+1) =V(t) + δ(t). Action values were calculated separately for each instrumental action (IPSI and CONTRA), all were set at 0 for the very first session, and for all subsequent sessions, V-values started at the final value of V for that action of the preceding session. The learning rate (α), was set at 0.3, however the pattern of data was preserved when learning rates were varied over a range of values (from 0.1-0.5) (**Fig S2C**).

There was a modification to this model that was made to separately assess the possibility of action values being updated on every press, which allowed for negative prediction errors in the absence of magazine entries, whereby δ was calculated to generate a negative term following all presses in which there wasn’t a rewarded magazine entry, and V was updated on every press.

Signal data was compiled across the Action window according to the value of V at the time of press: 0-0.3 (low); 0.3-0.6(low-med); 0.6-0.8(med-high) and 0.8-1(high). The same was done for δ during Outcome window: 0-.075(low); 0.075-0.15(low-med);0.15-0.23(med-high) and 0.23-0.3(high), with the maximum error term being equivalent to α.

### Video analysis

Video footage was analysed for key behavioral sessions using a markerless pose estimation approach. For experiment 1 (**Fig 3**), we analysed video data from the first and last sessions of contingency degradation training for 6 rats (12 videos, 2 per subject) and for experiment 2 (**Fig S5**) we analysed the mid-session reversal (6 videos, 1 per subject). Video footage was captured by Dahua cameras (DH-IPC-K35AP) situated centrally above the operant conditioning chambers (25 and 27 fps for experiments 1 and 2 respectively). Deeplabcut (Python 3.8.13, DLC version 2.2.1.1) was used to track the 2-D position of a number of body parts (nose, left ear, right ear, fiber optic cannula, neck, withers, rear, tail, contra paw, ipsi paw) on each frame for each video. Note that the video sets for the first and second experiments were processed and analysed separately, though the procedure was identical.

For each video set, 300 frames were labelled (25 or 50 frames per video for the first and second experiments respectively, extracted automatically via kmeans clustering) by a single experimenter and used to train a ResNet-50 neural network for 800,000 iterations. This trained network was then used to analyze each video, providing the inferred X-Y positions and likelihood values of each of the labelled body parts in each video frame. Inferred positions for a given body part with a likelihood value of less than 0.95 were discarded and then estimated via interpolation using a moving median with a window of 6 (*fillmissing* with *movmedian* in MATLAB).

Video tracking data was synchronized to behavioral data logged by MedPC at the start of each lever block by visually identifying the frames containing lever insertion, first lever press and lever retraction. A consistent feature of the environment above the magazine was used as the axis origin to provide a common space for videos recorded from different operant chambers. The neck was the midline feature that best provided reliably tracked points (above the likelihood cutoff) and so was used as a proxy for upper body position for the majority of video data analyses.

At the time of every lever press (onset as logged by MedPC) in each video, the inferred X-Y position data for each body part was determined and positional data specifically for the neck was extracted for one second before and one second after the lever press. The distance travelled by the neck between each frame was calculated in this 2-second peri-press period. These distance values were smoothed using a median filter over a window of 5 frames (*medfilt1* in MATLAB).

For each lever separately in each video, the median X-Y coordinates of all body parts at the time of press (position) as well as the median X-Y coordinates of the neck (trajectory) and the median distance travelled by the neck (distance) at each timepoint over the 2s peri-press period were calculated. For each subject, within-lever difference scores were calculated on each of these median datasets as either D1-D10 for degradation training or pre-post for the reversal session. To quantify whether basic features of movement surrounding the lever press significantly changed within subjects across degradation or reversal, we employed the same bCI procedure described above on the difference scores for trajectory (X and Y coordinates separately) and distance travelled. CIs that did not contain zero were identified, with a consecutive timepoint threshold corresponding to ∼0.25s used for determining significance (matching the consecutive threshold period used in the analysis of the dLight signal data).

### Statistical Analysis

For all analyses, non-orthogonal comparisons controlled the Type I error rate at α/k, for k contrasts, with α=0.05, according to the Bonferroni correction. For planned orthogonal comparisons, significance was set at α=0.05.

Behavioral data were analyzed according to planned contrasts comparing response rates on the degraded (or to-be-degraded) and non-degraded lever across instrumental training, contingency degradation training, and on test. Instrumental response rates during contingency degradation were converted into baseline scores, in order to account for pre-existing differences in response rates on each lever. This was taken as the response rate on each lever represented as a percentage of the mean response rate on that same lever across the final block of instrumental training (i.e. Block 4, Days 8-10).

For reversal training, orthogonal contrasts tested for differences in responding on the IPSI and CONTRA levers across training, reversal and reversal re-training.

For correlation analyses between AUC and inter-press and inter-event intervals, correlations were analyzed using simple linear regression, with Bonferroni-corrected p-values adjusted according to p(adjusted) = p*k, (k=12) for 12 comparisons (4 training blocks x 3 press types each for IPSI and CONTRA). Significance was set at p<0.05.

Details of the bCI procedure used for dLight waveform analyses and of the multiple regression analyses are described above. For analysis of fiber photometry AUCs across training (all experiments), pairwise comparisons (t-tests) were planned to test the difference between IPSI and CONTRA presses according to press type, within each training block, and averaged across all training blocks. During entries, tests compared the mean AUC during rewarded and empty magazine entries that followed either IPSI or CONTRA presses, within each training block. For assessment of prediction-error modulation of dopamine, planned comparisons tested the magnitude of dopamine within each press type at the highest and lowest V or δ values. For contingency degradation, comparisons were planned to test the difference between degraded and non-degraded presses within each press type. For mid-session reversal, comparisons were planned to test the difference between IPSI and CONTRA presses, averaged across all press types, before and after reversal, and within each press type after reversal. For all experiments, significance was set at p<α/k (Bonferroni).

Analysis of video tracking data (difference scores) was conducted using the bCI procedure described above. Pairwise comparisons (t-tests) were planned to compare X and Y coordinate data for each tracked body part at the time of lever press (median value per subject) on D1 and D10.

For outcome devaluation, behavioral and fiber photometry data were analyzed using planned, orthogonal contrasts testing for a main effect of side (IPSI vs CONTRA) and a main effect of value (Valued vs Devalued), controlling α at 0.05.

### Data availability

All source data are published in Figshare https://figshare.com/articles/dataset/Striatal_dopamine_encodes_the_relationship_between_actions_and_rewards/19083647. dLight1.1 was obtained under material transfer agreement with AddGene. All other materials for fiber photometry recordings were obtained from Tucker-Davis Technologies and Doric Lenses.

### Code availability

Custom MATLAB scripts are available upon request from Corresponding Author.

## Acknowledgments

The authors thank Amir Dezfouli for helpful comments on the manuscript, Myles Billard from Tucker-Davis Technologies for technical assistance with the fiber photometry recordings, and Karly Turner for assistance in the lab.

## Author contributions

Conceptualization: GH, TJB, CRN, BWB Methodology: GH, TJB, CRN, BWB Investigation: GH, TJB Funding acquisition: BWB Writing: GH, TJB, CRN, BWB

## Materials and Correspondence

All correspondence and materials requests should be directed to Bernard Balleine.

