## Supplementary figures for "Striatal dopamine encodes the relationship between actions and reward"

### Supplementary Figures S1 - S5

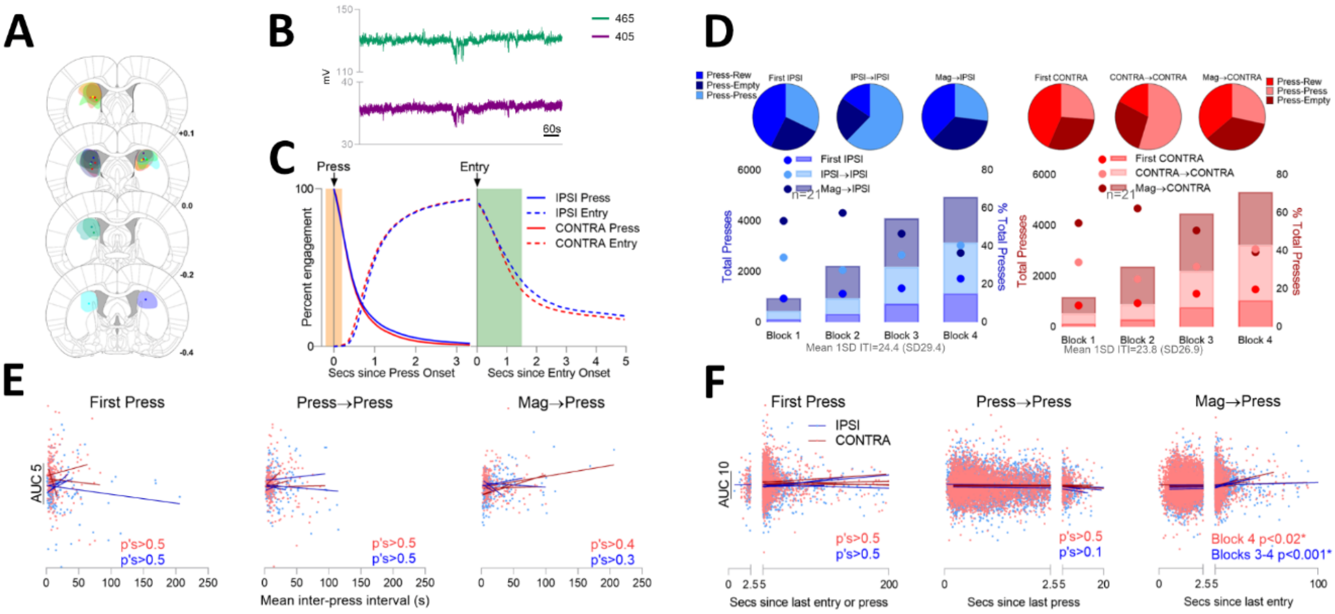


**Fig. S1. Related to Fig 1.** (**A**) Placement of fiber photometry cannula tips and spread of dLight1.1 in the pDMS for all rats included in the analysis. Numbers indicate anterior/posterior co-ordinates (mm) from bregma. (**B**) Representative demodulated fiber photometry traces for the 465 (dopamine binding dependent signal) and 405 (isosbestic control signal) excitation wavelengths. (**C**) Temporal distribution of engagement in behaviour for all logged behavioural events during training. Percent of animals engaged in IPSI (blue) and CONTRA (red) lever presses (solid lines) and magazine entries (dashed lines) in the period following the initiation (grey lines) of a lever press (left) or a magazine entry (right) Action window shaded in orange, Outcome window shaded in green. (**D**) Top: pie charts showing the percentage of each IPSI (left, blues) and CONTRA (right, reds) lever press subtype that were followed by either another press, an empty magazine entry, or a rewarded entry for all of instrumental training. Bottom: Total number of each lever press subtype within each training block across instrumental training (bars; left y-axis) and the percentage of total lever presses in that block for each press type (dots; right y-axis). (**E**) Mean (session-average) AUC (y-axis) during IPSI (blue) and CONTRA (red) lever press subtypes against the mean inter-press-interval for that session (x-axis). Each dot represents one training session. Lines indicate the best-fit slopes (simple linear regression). There were no significant correlations between press rate and dopamine release in any press type in any session, p-values indicated. (**F**) AUC (y-axis) during IPSI (blue) and CONTRA (red) press subtypes for each lever press against the inter-event interval since the preceding magazine entry or press (x-axis). Each dot represents one lever press event. Lines indicate the best-fit slopes (simple linear regression). There was no correlation between dopamine release and inter-event interval for IPSI or CONTRA First Presses or Press→Presses (p’s>0.1) however there was a positive relationship between both IPSI and CONTRA Mag→Presses and the time since their preceding magazine entry in the final blocks of training, indicating greater dopamine release for both IPSI and CONTRA lever presses with longer entry→press latencies, p-values indicated.

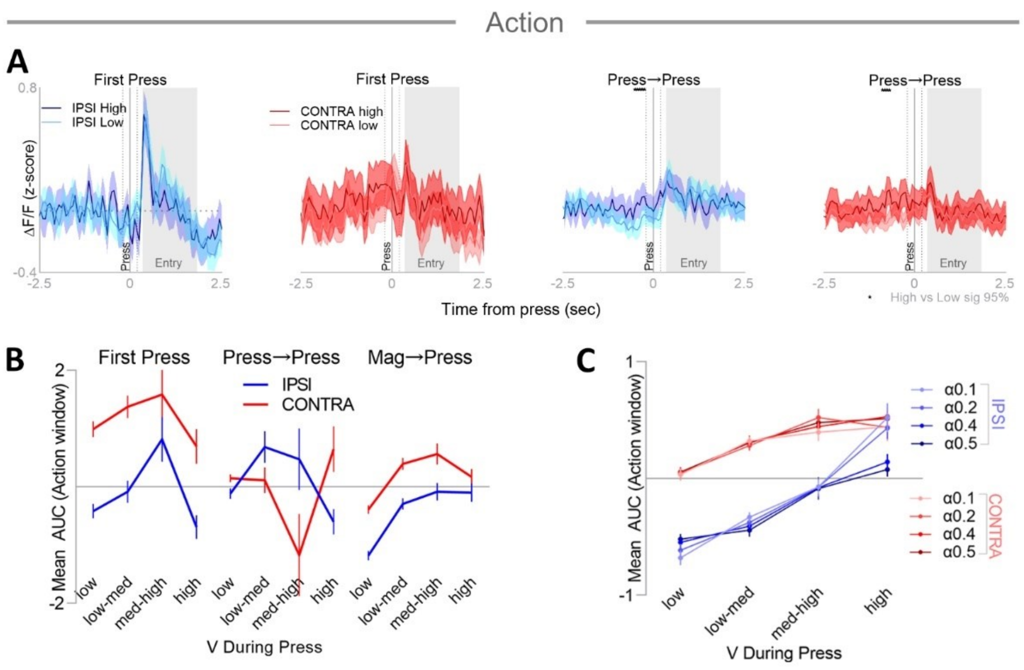


**Fig. S2. Related to Figure 2.** (**A**) Mean (±95% bCI) z-scored ΔF/F signals aligned to IPSI (blues) and CONTRA (reds) First Presses (left) and Press→Presses (right) with high (dark colors) or low (light colors) V-values. Asterisks indicate significant high vs low differences (95% bCI). (**B**) Mean (±SEM) AUC of the z-scored ΔF/F signal during the Action window for IPSI (blue) and CONTRA (red) lever press subtypes, subdivided according to action value when actions values are updated on every press. (**C**) Mean (±SEM) AUC during the Action window for all IPSI (blues) and CONTRA (reds) lever presses during instrumental training, subdivided according to action values, for varying alpha values (0.1-0.5).

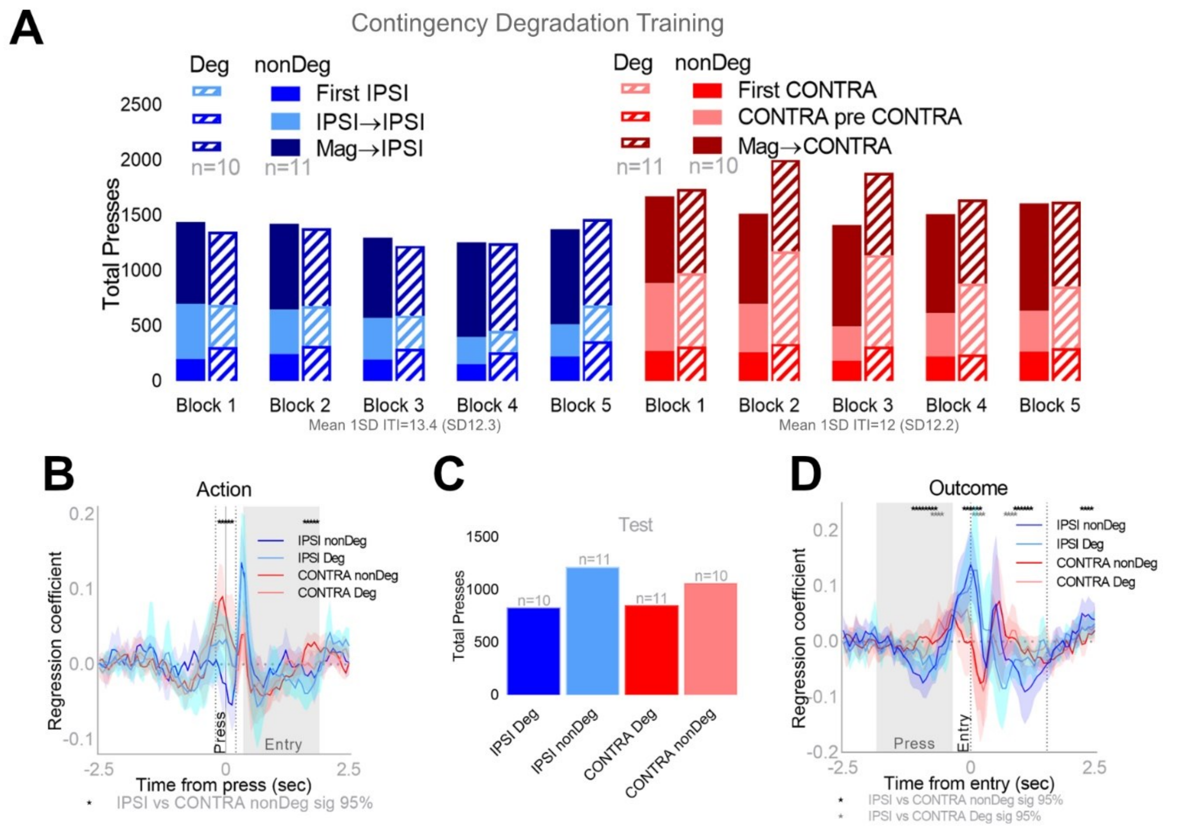


**Fig. S3. Related to Figure 4.** (**A**) Total number of each lever press subtype within each training block across contingency degradation training. (**B**) Mean regression coefficients (±95% bCI) for dLight signals aligned to IPSI (reds) and CONTRA (blues) lever presses that were nonDegraded (dark colors) and Degraded (light colors). Black asterisks indicate significant IPSI vs CONTRA nonDeg differences, grey asterisks indicate significant IPSI vs CONTRA Deg differences (95% bCI). (**C**) Total presses on each lever during the extinction test. **(D)** As for (B), aligned to magazine entries.

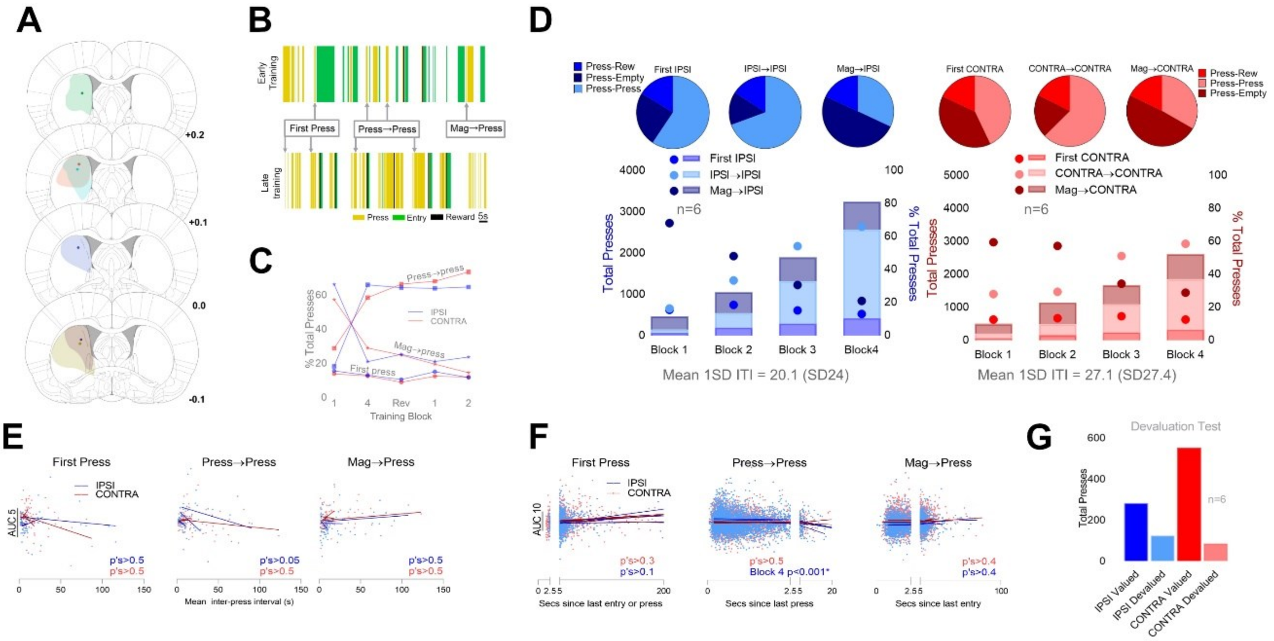


**Fig. S4. Related to Fig 5.** (**A**) Placement of fiber photometry cannula tips and spread of dLight1.1 in the pDMS for all rats included in the analysis. Numbers indicate anterior/posterior co-ordinates (mm) from bregma. (**B**) Representative sequence of events over 2 min for one rat during Block 1 of instrumental training (top) and the same rat in Block 4 of training (bottom). Lever presses in yellow, magazine entries in green and rewards in black. Example of lever press types identified. (**C**) Each lever press type represented as a percentage of total presses on the first and last blocks of instrumental training (blocks 1 and 4) and reversal training (blocks 1 and 2), and on the day of reversal. (**D**) Top: pie charts showing the percentage of each IPSI (left, blues) and CONTRA (right, reds) lever press subtype that were followed by either another press, an empty magazine entry, or a rewarded entry for all of instrumental training. Bottom: Total number of each lever press subtype within each training block across instrumental training (bars; left y-axis) and the percentage of total presses in that block for each press type (dots; right y-axis). (**E**) Mean (session-average) AUC (y-axis) during IPSI (blue) and CONTRA (red) lever press subtypes against the mean inter-press-interval for that session (x-axis). Each dot represents one session. Lines indicate the best-fit slopes (simple linear regression). There were no significant correlations between lever press rate and dopamine release for any press subtype in any session, p-values indicated. (**F**) AUC (y-axis) during IPSI (blue) and CONTRA (red) lever press subtypes, against the inter-event interval since the preceding magazine entry or press (x-axis), for every press. Each dot represents one lever press. Lines indicate the best-fit slopes (simple linear regression), p-values indicated. (**G**) Total presses on each lever during the devaluation test.

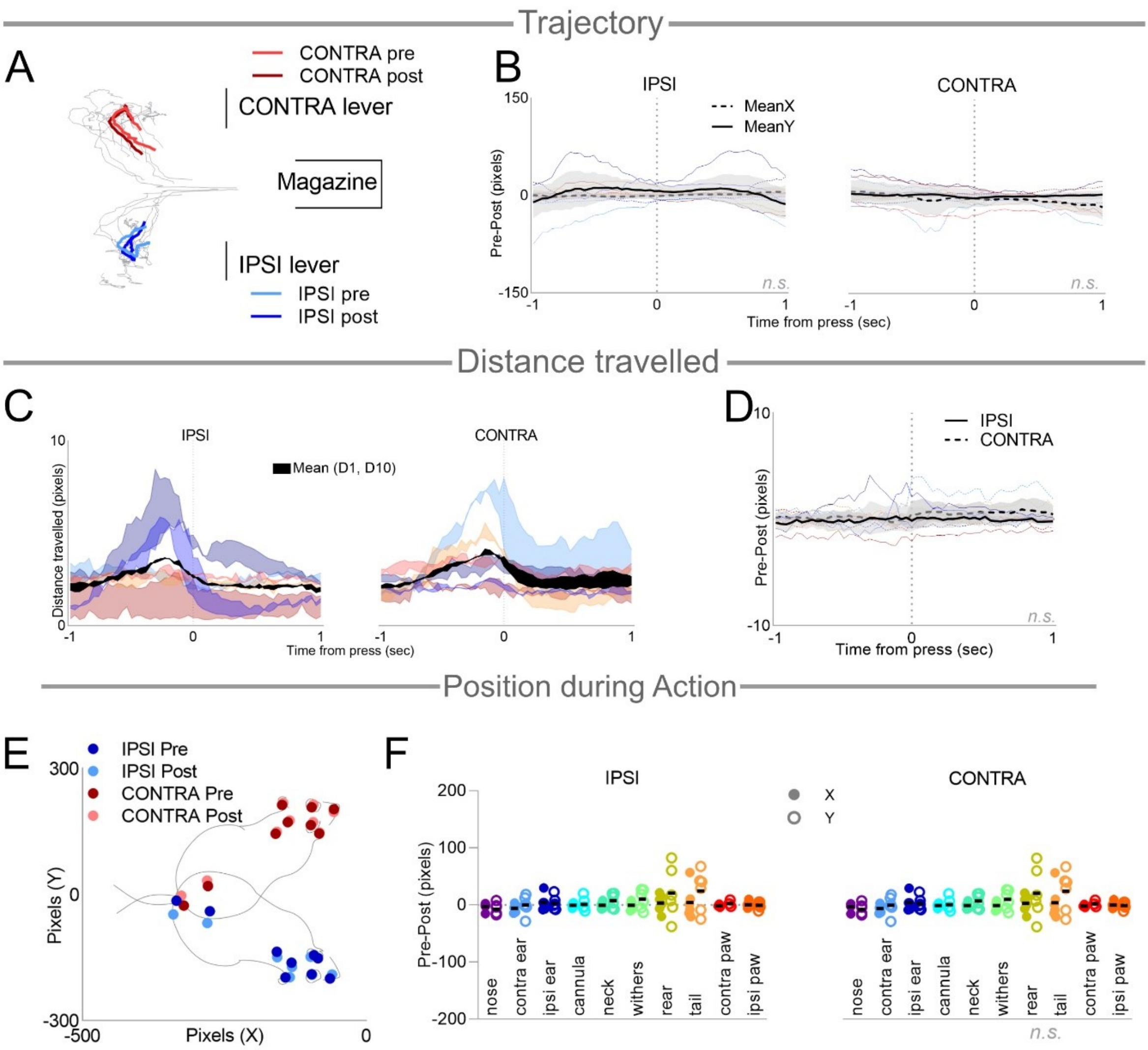


**Fig. S5. Related to Figure 5.** (**A**) The mean trajectory of all animals on each lever across the 2-s peri-press period (colored lines), before reversal (pre, light colors) and after reversal (post, dark colors). Median trajectory for each individual animal shown in grey. (**B**) Difference scores (Pre-Post) of co-ordinates in (A), X co-ordinates in dotted lines, Y co-ordinates in solid lines. The mean (±95% bCI) difference score of all animals is shown in black, and the difference scores of each individual animal are in colored lines. (**C**) Mean distance travelled in the 2-s peri-press period per frame pre- and post-reversal on the IPSI (left) and CONTRA (right) levers, with shading between the two lines to indicate the difference. The mean for all animals is represented in black, with the median for each rat plotted in the coloured lines for pre and post reversal. (**D**) Difference scores (Pre-Post) for lines represented in (C). The mean (±95% bCI) difference score of all animals is shown in black (IPSI solid lines, CONTRA dotted lines), and the difference scores of each individual animal are in colored lines. (**E**) The mean X-Y position of 10 tracked body parts for all rats at the time of press on the IPSI (blues) and CONTRA (reds) levers. Pre reversal (dark dots) and post reversal (light dots). (**F**) Difference scores along X (filled circles) and Y (open circles) co-ordinates for each marker for each rat, mean indicated by black bar.
